# The mitochondrial cohibitin complex facilitates the biogenesis of inner membrane proteins

**DOI:** 10.64898/2026.01.12.699002

**Authors:** Büsra Kizmaz, Annika Nutz, Steffen Hess, Johannes Hagen, Sergio Guerrero-Castillo, Alfredo Cabrera-Orefice, Chang-Mo Yoo, Christof Osman, Markus Räschle, Hyun-Woo Rhee, Johannes M. Herrmann

## Abstract

The inner membrane of mitochondria contains many membrane-embedded carrier proteins of the SLC25 family to facilitate the exchange of metabolites between the cytosol and mitochondria. These carriers use a specific import route for their biogenesis that relies on the TIM22 complex as an inner membrane translocase. The molecular details of carrier biogenesis are not well understood. Using an improved, desthiobiotin-based proximity labeling approach called Destni in living yeast cells, we identified the mitochondrial protein Aim11 as an interactor of newly imported carrier proteins. Aim11 forms a 160 kDa complex together with Iai11, Gep7 and Mtc1 in the mitochondrial inner membrane that we named the comrade-of-prohibitin (cohibitin) complex owing to its genetic interaction with prohibitins. Deletion of Aim11 impairs the import of carrier proteins into the inner membrane and renders cells hypersensitive to carrier overexpression. Our data suggest that the cohibitin complex plays a quality control function that supports the TIM22-mediated insertion of carrier proteins into the inner membrane of mitochondria.

**Summary:** Kizmaz et al. identified Aim11 as a novel quality control factor that facilitates the insertion of carrier proteins into the inner membrane of mitochondria. Aim11 is part of the membrane-embedded cohibitin complex which cooperates with prohibitins in inner membrane protein biogenesis.

## Introduction

Mitochondria are essential organelles of eukaryotic cells that play central roles in metabolism, energy production, and signaling. Mitochondria contain their own genome that codes for a small set of hydrophobic proteins (13 in humans and 8 in baker’s yeast) that are part of the respiratory chain and the ATP synthase complex (Ott et al., 2016). All other proteins (about 1,500 in humans and 900 in baker’s yeast) are nuclear encoded and synthesized in the cytosol (Morgenstern et al., 2017; Rath et al., 2021). Targeting signals in these ‘precursor’ proteins direct them to the mitochondrial surface, where they are bound by receptor proteins. Protein translocases in the mitochondrial outer and inner membranes then import the precursors into mitochondria and sort them to their appropriate intramitochondrial location (Wiedemann and Pfanner, 2017). Mitochondria of animal and fungal cells contain three protein translocases: The translocase of the outer mitochondrial membrane (TOM) complex forms large pores in the outer membrane that serve as a general entry gate for proteins (Araiso et al., 2022). Two translocases of the inner membrane (TIM23 and TIM22 complexes) mediate the translocation into the matrix and the insertion of inner membrane proteins (Jain et al., 2025; Kang et al., 2018; Kizmaz et al., 2024; Wiedemann and Pfanner, 2017). The TIM complexes do not form aqueous pores but rather protein-lipid interphases to facilitate the translocation of proteins across the inner membrane (Fielden et al., 2023; Sim et al., 2023). The TIM23 complex, which is coupled to the ATP-driven import motor serves precursors with N-terminal presequences that act as matrix-targeting signals (MTSs)(Mokranjac, 2020). Clients of the TIM22 complex lack N-terminal presequences and mostly expose their N-termini to the intermembrane space (IMS). Many TIM22 substrates are members of the solute carrier (SLC25) family (Koehler et al., 1998; Nauerz et al., 2025). These ‘carriers’ (35 in yeast and 53 in humans) facilitate the exchange of metabolites, nucleotides and metals between mitochondria and the cytosol (Horten et al., 2020; Kang et al., 2018). Whereas the import into the matrix by the TIM23 complex is well studied, the biogenesis of carriers via the TIM22 complex-dependent pathway still awaits a detailed molecular characterization.

Protein translocation into mitochondria is under surveillance of protein quality control systems which remove stalled translocation intermediates (Song et al., 2021). Ubiquitin ligases are positioned at the cytosolic site of the TOM complex to promote proteasomal degradation of non-productive import intermediates (Mårtensson et al., 2019; Schulte et al., 2023). In addition, proteases of the inner membrane expose proteolytic domains to the matrix and the IMS (m-AAA and i-AAA protease, respectively) to recognize and remove defective or orphaned newly imported proteins or to adapt the composition of the import machinery to prevailing conditions (Deshwal et al., 2020; Elancheliyan et al., 2024; Hsu et al., 2025).

The prohibitin complex of the inner membrane supports the folding and assembly of membrane proteins and, if this is not possible, facilitates their degradation (Steglich et al., 1999). This ring-shaped complex of alternating prohibitin 1 and 2 (Phb1, Phb2) subunits organizes a bell-shaped scaffold in the inner membrane that is associated with the AAA proteases (Kohler et al., 2023; Lange et al., 2025; Nijtmans et al., 2002; Osman et al., 2009b; Tatsuta and Langer, 2017; Wei et al., 2017). The structure and function of the prohibitin complex is presumably ubiquitously conserved in mitochondria and bacteria, where it is referred to as the FtsH-HflKC complex (Gao et al., 2025; Ghanbarpour et al., 2025; Hong et al., 2025; Lange et al., 2025; Qiao et al., 2022). Despite its conservation from bacteria to eukaryotes and its relevance in the context of physiology and diseases (see (Tatsuta and Langer, 2017) for overview), the molecular function of the prohibitin complex is only poorly understood. Recently, an even larger structure was identified in the inner membrane, called the MIMAS complex which functions as a biogenesis and metabolic platform, consisting of a multitude of different proteins (Horten et al., 2024; Moretti-Horten et al., 2024). MIMAS is distinct from the prohibitin complex and its specific role in membrane protein biogenesis is not clear.

Quality control factors which support the insertion, topogenesis and folding of carrier proteins in the mitochondrial inner membrane have not been identified so far. A role for carrier biogenesis was neither reported for prohibitins nor for the MIMAS complex. We therefore wondered whether newly imported carrier proteins interact with other, so far unidentified proteins of the mitochondrial inner membrane. To identify such factors, we employed a proximity labeling approach with an optimized version of a biotin ligase fused to the N-terminus of the oxodicarboxylate carrier Odc1. We identified the inner membrane protein Aim11 as a potential interactor of newly imported carriers. The direct interaction of carriers and Aim11 was confirmed by pull-down experiments with newly imported carrier proteins.

Aim11 is part of a complex of 160 kDa which we named the cohibitin complex. Aim11 and the other cohibitin subunits show synthetic growth defects with Phb1 deletions, suggesting that cohibitin and prohibitin complexes might function in parallel, synergistic pathways. Our study identifies Aim11 and the cohibitin complex as novel factors that facilitate the efficient biogenesis of inner membrane proteins of mitochondria.

## Results

### Proximity labeling identifies Aim11 as an Odc1 interactor

The biogenesis of carrier proteins was mainly studied by *in vitro* import reactions with isolated mitochondria. These import experiments allowed the definition of distinct stages of the import reaction (Pfanner and Neupert, 1987). However, the final step by which the carriers are inserted into the inner membrane and folded into their three-dimensional structure is still poorly understood. The structures of several carrier proteins have been solved by crystallography (Jones et al., 2023; Ruprecht et al., 2014; Ruprecht et al., 2019), confirming closely the structures proposed by AlphaFold (Abramson et al., 2024; Jumper et al., 2021).

Carriers resemble membrane-embedded cylinders formed by three structurally conserved carrier ‘modules’ (Fig. 1A). Each module consists of a hairpin-like arrangement of two transmembrane domains and a short matrix loop. Whether each module is recognized and inserted sequentially by TIM22 so that carriers fold only after their complete insertion into the membrane, or whether carrier folding is facilitated on the TIM22 is unknown (Fig. 1B). Each module has been shown to contain sufficient information for mitochondrial targeting compatible with a sequential insertion model (Brandner et al., 2005; Wiedemann et al., 2001).

**Figure 1.**
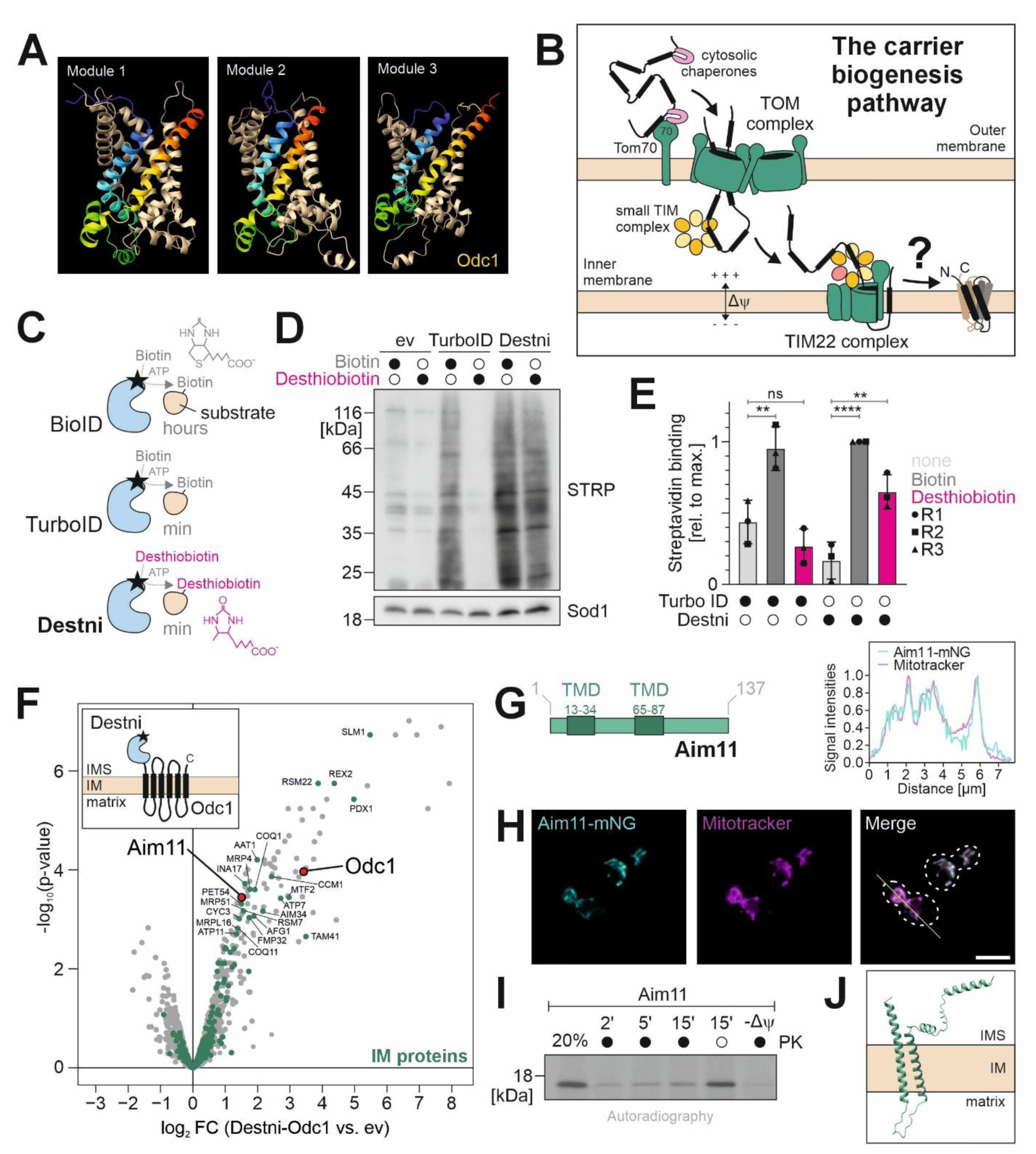
Proximity labeling identifies Aim11 as an Odc1 interactor. **(A)** The structure of the carrier Odc1 of *S. cerevisiae* was predicted using AlphaFold 3 (Abramson et al., 2024). The three modules are highlighted in rainbow coloring to show the repetitive arrangement of the protein structure which is characteristic for members of the SLC25 carrier family (Ruprecht et al., 2019). **(B)** Schematic representation of the carrier import pathway. **(C)** Schematic representation of the (desthio)biotinylation reactions catalyzed by optimized versions of the biotin ligase BirA (Branon et al., 2018; Roux et al., 2012). **(D and E)** Wild type cells were inoculated in low-biotin medium, shifted to medium without biotin overnight, and Destni and TurboID were expressed in the cytosol for 4 h. The labeling reaction was started by adding 100 µM biotin or desthiobiotin. Cells were harvested, lysed, and analyzed via Western blot. Levels of biotinylated proteins were detected with horse radish peroxidase-coupled streptavidin (STRP). Sod1 served as a loading control. (Right) Whole lane signals from three biological (independent) replicates were quantified. Shown are mean values and standard deviations. Statistical analysis was performed using a two-way ANOVA with Sidak’s multiple comparison test. Statistical significance was assigned as follows: p-value < 0.01 = **; < 0.0001 = ****. ns, not significant. **(F)** Volcano plot showing the interactome comparison of the Destni-fused carrier Odc1 with a control for three biological replicates. Proteins were eluted with biotin from the streptavidin beads. Mitochondrial inner membrane proteins are depicted in green. The bait protein Odc1 and Aim11, a potential interactor of Odc1, are highlighted in red. log_2_ FC, log_2_ fold change. (Insert) Schematic representation of the fusion protein comprised of the N-terminal desthiobiotin ligase Destni with Odc1, a mitochondrial oxodicarboxylate carrier. See also Table S3 for the dataset. **(G)** Schematic diagram of the Aim11 sequence highlighting the membrane-spanning regions. TMD, transmembrane domain. **(H)** Genomically tagged Aim11-mNeonGreen (Aim11-mNG) strains were grown in full galactose medium to mid-logarithmic phase and visualized by fluorescence microscopy. Shown are the channels for mNeonGreen, Mitotracker Orange to image the mitochondrial network, and the merge. Scale bar: 5 µm. (Top) Min-max normalized plot profiles of the fluorescent signals along the yellow line shown in the merge. (**I**) Radiolabeled Aim11 was incubated with isolated mitochondria from wild type cells. Samples were incubated at 30°C for the indicated time points. Non-imported protein was removed by proteinase K (PK) treatment. A control sample was treated with a mix of valinomycin, oligomycin, and antimycin A to dissipate the membrane potential (-Δψ). The 20% sample shows one-fifth of the radiolabeled protein used per time point of the import reaction. (**J**) Schematic topology of Aim11 based on the AlphaFold 3 structure (Abramson et al., 2024).

We followed a proximity labeling approach to identify interactors of inserting carrier proteins in living yeast cells. The BirA biotin ligase of *E. coli* has been optimized by engineering steps to compromise its native, mono-specific substrate specificity so that it biotinylates all proteins unambiguously within a certain distance (BioID) and to increase its activity to speed up the biotinylation (TurboID) (Branon et al., 2018; Roux et al., 2012). However, the application of TurboID in yeast has important limitations, as efficient labeling typically requires biotin supplementation, which can stimulate endogenous biotinylation reactions and perturb the metabolic landscape of the organism (Fenech et al., 2023). We therefore employed Destni (for desthiobiotin ligase), a TurboID-derived ligase engineered by yeast surface display to efficiently utilize desthiobiotin as an alternative substrate (Fig. 1C, S1A, B). Because yeast cells do not contain endogenously desthiobiotinylated proteins, Destni enables proximity labeling using desthiobiotin without inducing the widespread endogenous biotinylation associated with biotin supplementation. Expressing Destni in the cytosol resulted in a quick (desthio)biotinylation of proteins, reaching maximal levels already after 30 minutes (Fig. 1D, E, S1C-E). Whereas TurboID conjugated proteins only with biotin, the Destni-catalyzed reaction also resulted in an efficient desthiobiotinylation of proteins.

Next, we fused the TurboID and Destni domains to the N-termini of three mitochondrial carrier proteins (Pet9, Odc1, Mir1) as carriers tolerate N-terminal but no C-terminal extensions (Breker et al., 2013). Pet9 represents the highly abundant ATP/ADP carrier, Odc1 transports oxodicarboxylates and Mir1 phosphate. All fusion proteins were expressed in wild type cells to catalyze the (desthio)biotinylation of proteins (Fig. S1F-H). We depleted the endogenous biotin levels in biotin-free medium, induced the expression of the Destni-carrier fusion proteins, added desthiobiotin, lysed the cells and purified desthiobiotinylated proteins (Fig. S1I). The Destni-Odc1 protein was found to be efficiently self-desthiobiotinylated, whereas the other two carrier fusion proteins were only moderately enriched in comparison to the controls (Fig. S1J, Table S3). We therefore focused on the Destni-Odc1 samples which pulled down many mitochondrial inner membrane proteins consistent with the mitochondrial location of the protein (Fig. S1K). Several poorly or non-characterized proteins were found among the consistently desthiobiotinylated proteins in the Destni-Odc1 samples, including the inner membrane protein Aim11 (Fig. 1F). Since this protein had not been characterized before, we decided to further study the potential connection of Aim11 to carrier biogenesis.

### Aim11 is a mitochondrial inner membrane protein

Aim11 is a short protein of 137 residues with two predicted transmembrane domains (Fig. 1G). Aim11 was reported before as a protein of the mitochondrial proteome (Morgenstern et al., 2017). We confirmed the mitochondrial location of Aim11 by fluorescence microscopy with strains expressing an Aim11-mNeonGreen fusion protein; the signal of the mitochondria-staining dye mitotracker colocalized with that of Aim11 (Fig. 1H).

Aim11 lacks an N-terminal MTS (its TargetP score is 0.03) but contains an internal MTS (iMTS) sequence around residue 20 that is adjacent to the first of its two predicted transmembrane domains (Fig. S2A) (Emanuelsson et al., 2007; Jung et al., 2024). Such patterns were found in mitochondrial inner membrane proteins that insert in a loop-like structure and expose their N termini to the IMS (Fölsch et al., 1996). To elucidate the import properties in Aim11, we synthesized radiolabeled Aim11 in reticulocyte lysate and incubated the protein with isolated mitochondria before non-imported protein was removed by treatment with proteinase K (Fig. 1I). In the presence of ATP and NADH, Aim11 was imported and reached a protease-protected location, but Aim11 was not imported when the inner membrane potential was dissipated. The import of Aim11 into mitochondria of temperature-sensitive *tim17-5* was reduced (Fig. S2B), whereas the import into *tim22-14* mitochondria occurred largely unaffected (Fig. S2C). The imported Aim11 became accessible to proteinase K when the outer membrane of mitochondria was ruptured by hypotonic swelling (Fig. S2D) and was not extractable with carbonate (Fig. S2E). Thus, Aim11 represents an inner membrane protein of mitochondria that reaches its location via the TIM23 pathway, using an internal targeting sequence, consistent with the topology proposed by AlphaFold 3 (Abramson et al., 2024) (Fig. 1J).

### Aim11 is part of an inner membrane complex

To identify potential interaction partners of Aim11, we expressed the protein with a C-terminal hemagglutinin (HA). This Aim11-HA protein complemented the moderate growth defect of the Δ*aim11* mutant (Fig. S3A) and was detectable in Western blots in whole cell extracts (Fig. S3B). We isolated mitochondria from Aim11-HA and control cells (three independent replicates each), lysed them with the mild detergent digitonin and incubated the extracts with anti-HA antibody-coupled magnetic beads. Aim11 and the three inner membrane proteins Gep7, Iai11, and Mtc3 were significantly enriched in the Aim11-HA pulldowns, suggesting that these proteins are complex partners (Fig. 2A). The sequence and structure of Iai11 is related to that of Aim11, and Gep7 and Mtc3 are also paralogs (Fig. 2B, S3C, D). In addition, Cox6, a subunit of cytochrome oxidase was recovered in the Aim11-HA fraction.

**Figure 2.**
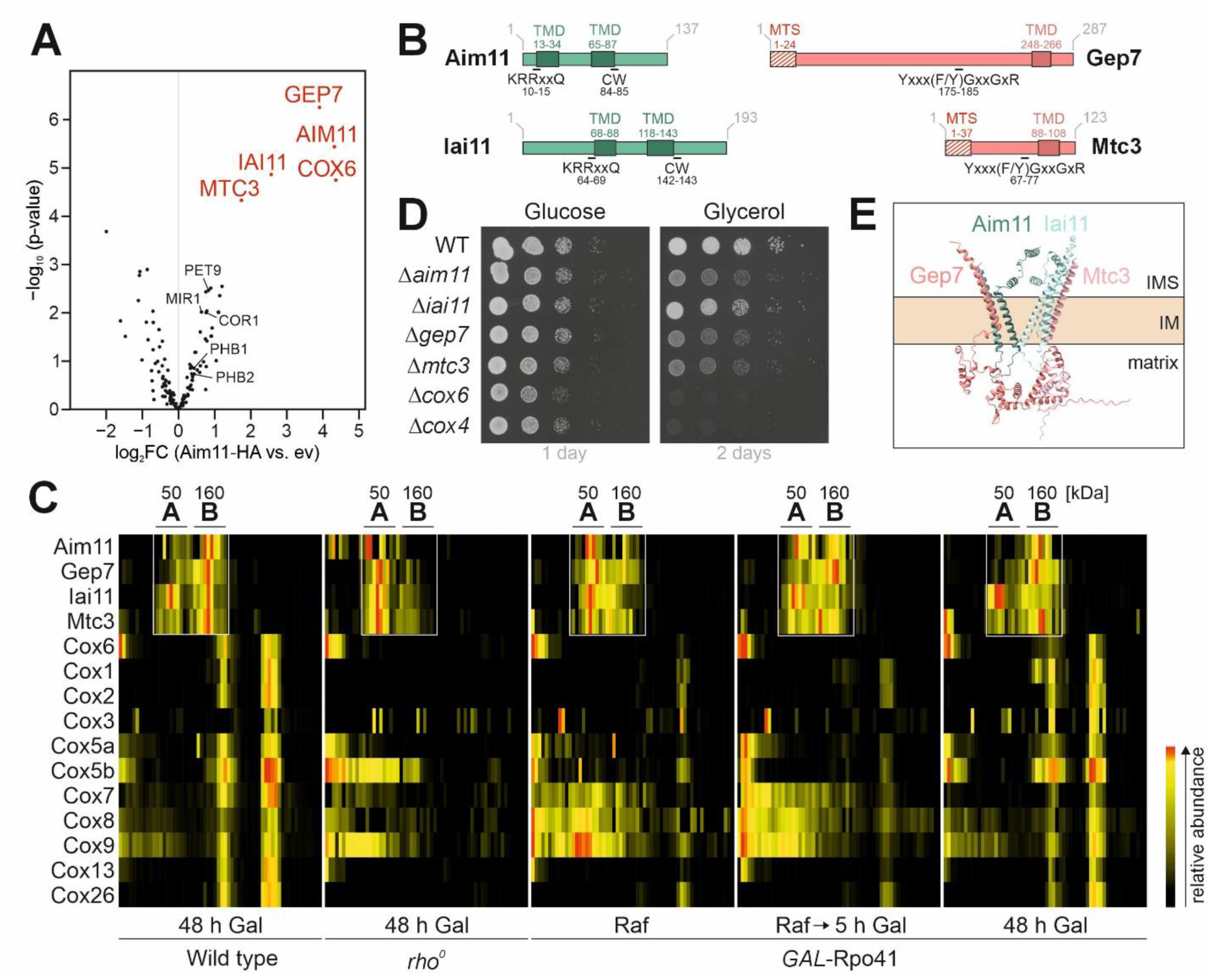
Aim11 is part of an inner membrane complex. **(A)** Crude isolated mitochondria of wild type cells containing an empty plasmid (ev) or an Aim11-HA expression plasmid were grown in selective galactose medium to mid-logarithmic phase and lysed with 1% (w/v) digitonin. The samples were subjected to affinity chromatography with anti-HA antibody-coupled magnetic beads. Bound proteins were analyzed by mass spectrometry. A volcano plot depicting the interactome of the indicated samples is shown. The most abundant proteins are labeled in red. log_2_ FC, log_2_ fold change. Please refer to Table S5 for the entire dataset. **(B)** Schematic representation of the structures of the indicated proteins. TMD, transmembrane domain. **(C)** Wild type, *rho^0^* and *GAL*-Rpo41 cells were grown in raffinose (Raf) or galactose (Gal) as indicated, lysed in digitonin buffer and subjected to BN-PAGE. Slices of the gels were cut out, and containing proteins were analyzed by proteomics. Please refer to Table S9 for the full dataset. **(D)** Wild type (WT) and the indicated deletion mutants were grown on galactose-containing media to mid-logarithmic phase before tenfold serial dilutions were dropped onto the indicated plates and incubated at 30°C. **(E)** AlphaFold 3 prediction of the Aim11-Gep7-Iai11-Mtc3 complex (Abramson et al., 2024).

To analyze the potential association of these mitochondrial proteins, we used complexome profiling that is based on co-migration on blue native polyacrylamide gels (BN-PAGE). To assess whether the mitochondrial genome is critical for the complex size of these proteins, we analyzed wild type and *rho^0^* cells (i.e. cells lacking mitochondrial DNA) as well as mutants in which the expression of the mitochondrial RNA polymerase Rpo41 was under control of the *GAL10* promoter (*GAL*-Rpo41). When cells were grown on raffinose medium, mitochondrial transcription was repressed in these strains. When cells were shifted to galactose medium for different time periods, mitochondrially encoded genes were again expressed and accumulated over time. Mitochondria were isolated from these strains, lysed with digitonin and subjected to BN-PAGE (4-16% acrylamide concentration, Fig. S4A-C). Each lane was cut into 60 slices, which were individually analyzed by tandem mass spectrometry. Aim11, Gep7, Iai11, and Mtc3 co-migrated in a range between 50 and 160 kDa, forming two maxima. In wild type mitochondria, the proteins mainly populated the 160 kDa complex, whereas in the absence of mitochondrial DNA or RNA, the 50 kDa species was more abundant (Fig. 2C). Both species were clearly distinct from the cytochrome oxidase, which co-migrated with Cox6 (Fig. 2C, S4C). Moreover, the phenotypes of deletion mutants of the five proteins confirmed the clear difference between Cox6 and the other four proteins. Cox6 was essential for respiration (such as the cytochrome oxidase protein Cox4), whereas the deletion of Aim11, Gep7, Iai11 and Mtc3 resulted in a milder growth defect on respiratory carbon sources (Fig. 2D).

Next, we used AlphaFold 3 (Abramson et al., 2024) to model the structure of Aim11-Gep7-Iai11-Mtc3 as well as (Aim11)_2_-(Gep7)_2_ complexes (Fig. 2E, S3E). Both complexes were predicted to constitute V-shaped helix bundles that are deeply embedded into the inner membrane with a wide opening towards the IMS. Our attempts to confirm this organization by crosslinking combined with mass spectrometry failed due to very low crosslinking efficiency between these membrane-embedded subunits. In summary, Aim11 is part of a heterooligomeric complex that contains the proteins Gep7, Iai11 and Mtc3, and potentially a small subfraction of Cox6.

### The Aim11 complex is essential in prohibitin-deficient cells

The function of Aim11, Gep7, Iai11 and Mtc3 have not been characterized so far, but their common presence in one complex has been reported before (Morgenstern et al., 2017). All four proteins were identified in the proteome of highly purified mitochondria (Morgenstern et al., 2017) and we confirmed the mitochondrial location by the expression of mNeonGreen fusion proteins (Fig. 1H, S3F). Surprisingly, all four proteins, as well as Cox6 had been identified in a screen for components that become essential upon deletion of the prohibitin protein Phb1 (Osman et al., 2009a). In addition to these five proteins, several biogenesis factors of inner membrane proteins were found among the 35 hits of the synthetic lethality screen with *phb1* deletion mutants, including Oxa1, Atp10, Atp23, Mdm10, Ups1, Yta10, and Yta12.

To confirm the genetic interaction with the prohibitin-encoding genes, we crossed single deletion mutants of *AIM11* and *GEP7*, as well as *OXA1* for control, in a Mat *a* background with strains lacking *PHB1* in Mat *alpha* cells. From the diploid progeny, spores were derived by meiosis and the tetrads were analyzed. Drop dilution experiments confirmed the synthetic growth defect of *Δaim11Δphb1* and *Δgep7Δphb1* double mutants (Fig. 3A, B) as well as of *Δoxa1Δphb1* cells (Fig. S3G). Owing to the profound co-dependence of the prohibitins with the Aim11 complex, we named the Aim11/Gep7/Iai11/Mtc3 complex cohibitin complex for comrade-of-prohibitin.

**Figure 3.**
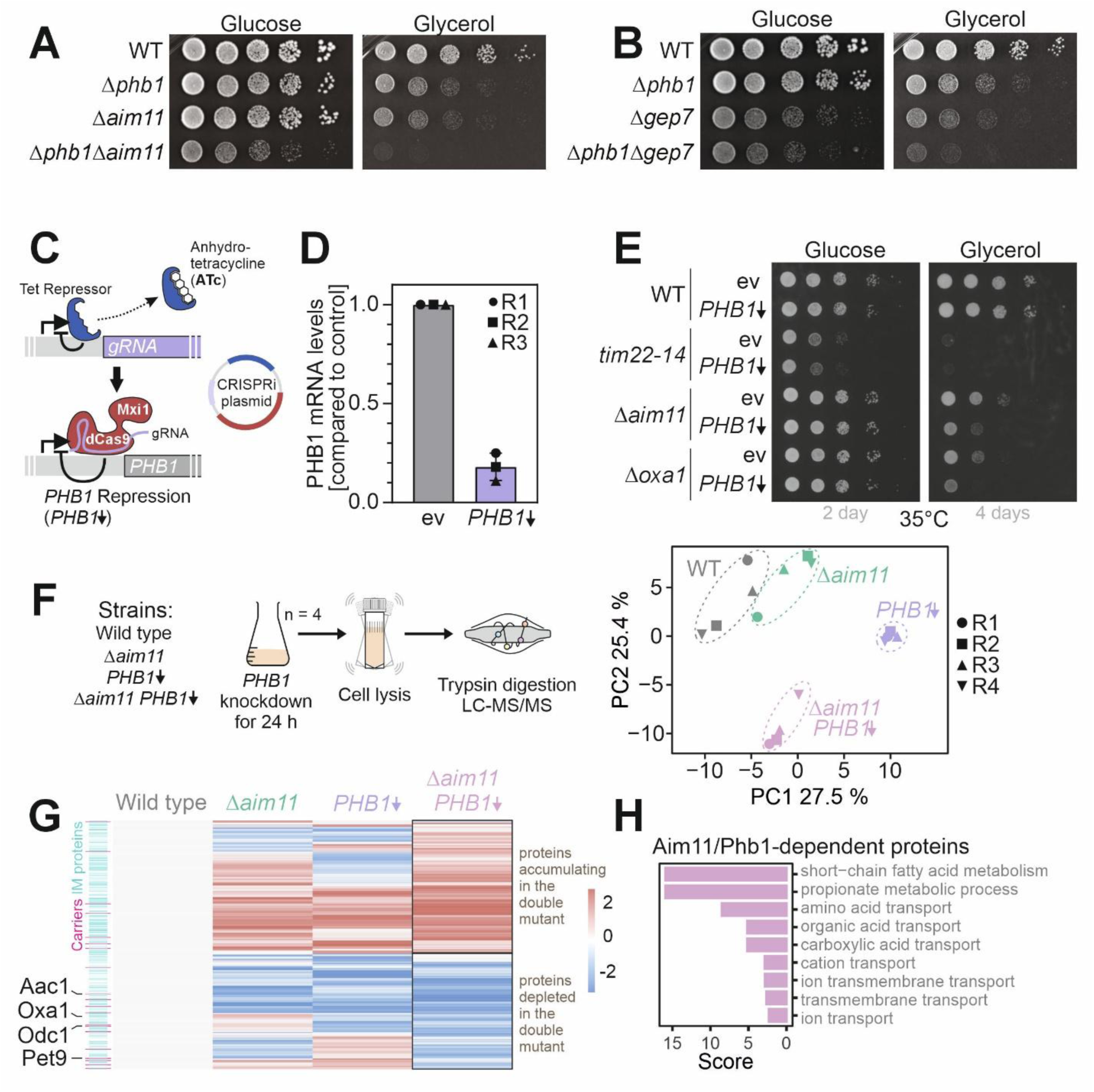
*AIM11* exhibits a negative genetic interaction with *PHB1*. (A and. **B)** Wild type (WT) and the indicated deletion mutants were grown on galactose-containing media to mid-logarithmic phase before tenfold serial dilutions were dropped onto the indicated plates and incubated at 30°C. **(C)** Schematic representation of the CRISPRi approach. The Tet repressor is inhibited by anhydrotetracycline (ATc), which leads to the expression of a specific guide RNA (gRNA). The gRNA directs the dCas9-Mxi1 fusion to the promoter of a gene of interest, thereby blocking transcription of the respective gene, which is *PHB1* in this case. **(D)** A CRISPRi plasmid containing a sequence for gRNA against *PHB1* or without any gRNA sequence (empty vector, ev) was transformed into wild type cells. Cells were grown to mid-logarithmic phase, and the knockdown of *PHB1* was induced with 960 ng/ml ATc. RNA was isolated after 2 h of induction, and mRNA levels were analyzed by RT-qPCR. The mean values and standard deviations of three independent replicates (n = 3) are shown. **(E)** The indicated strains were grown in galactose medium to mid-logarithmic phase, and *PHB1* repression was induced with 960 ng/ml ATc. After 6 h, a ten-fold dilution series was dropped on agarose plates containing glucose or glycerol as carbon sources. Plates were incubated at 35°C for the indicated number of days. **(F)** (Left) Scheme for the analysis of *PHB1* knockdown by mass spectrometry. (Right) Principal component (PC) analysis shows the similarity of the protein spectra identified in the different samples. R, replicate. **(G)** Heatmap of the mitochondrial proteome data gained in the mass spectrometric analysis from F. Z-scores were calculated using the mean of the log_2_ fold changes of n = 4 biological replicates and normalized to the wild type signals. See also Table S7 for the output of hierarchical clustering. Inner membrane proteins are highlighted in cyan, carrier proteins in magenta. **(H)** Proteins that were explicitly diminished in the *Δaim11 PHB1↓* samples (log_2_ fold change > - 0.5 and p-value <0.05) were analyzed by gene ontology (GO) enrichment using the GOrilla tool (http://cbl-gorilla.cs.technion.ac.il).

What is the underlying basis of this co-dependence? To study this, we depleted Phb1 in *Δaim11* cells by RNA interference (Backes et al., 2021; Smith et al., 2016) (Fig. 3C) which reduced the levels of *PHB1* mRNAs by 80% (Fig. 3D) and confirmed the synthetic negative growth defect in *Δaim11* and *Δoxa1* cells (Fig. 3E). To better characterize the molecular defects underlying the negative genetic interaction, we depleted *PHB1* mRNAs for 24 h in wild type and *Δaim11* cells (Fig. 3F) and analyzed the whole cell proteomes by mass spectrometry. Principle component analysis of the protein patterns showed that the single

*Δaim11* deletion had a much smaller effect on the proteome than the loss of Phb1. However, the double deletion considerably differed from both single mutants, in line with the strong synthetic effect observed in respect to cell growth. The abundance of many mitochondrial proteins was changed in the double mutant (Fig. 3G). Apparently, the simultaneous loss of the Aim11 and Phb1 left a strong footprint on the mitochondrial proteome. Interestingly, many carrier proteins were found to be depleted in the double mutant, including Pet9 and Odc1. The unbiased analysis of the GO terms of proteins that were found to be depleted in the double mutant supported the relevance of Aim11 and Phb1 for the biogenesis of metabolite transporters, i.e. the GO terms that represent mitochondrial carriers (Fig. 3H). From this, we conclude that the prohibitin and cohibitin complexes play a partially redundant role in the biogenesis of mitochondrial inner membrane proteins, including the members of the mitochondrial carrier family.

### Carrier proteins interact with the complex during membrane insertion

Carrier proteins are inserted into the inner membrane of mitochondria by the conserved TIM22 complex (Fig. 4A). We wondered whether the cohibitin complex acts upstream, downstream or in parallel to the TIM22 complex. We therefore carried out *in vitro* import experiments with radiolabeled Pet9 carrier protein and mitochondria isolated from wild type and *Δaim11* cells. Whereas Pet9 was imported efficiently into wild type mitochondria so that it became protected against added proteinase K, only very low amounts of Pet9 were imported into *Δaim11* mitochondria (Fig. 4B, C). This import defect was specific for carrier proteins and only mildly affected matrix proteins such as Su9-DHFR (Fig. 4D, E). We therefore regard it as unlikely that the reduced carrier import is indirectly caused by a lower membrane potential in the *Δaim11* mitochondria, even though *Δaim11* cells have a somewhat reduced respiration activity (Fig. 4F).

**Figure 4.**
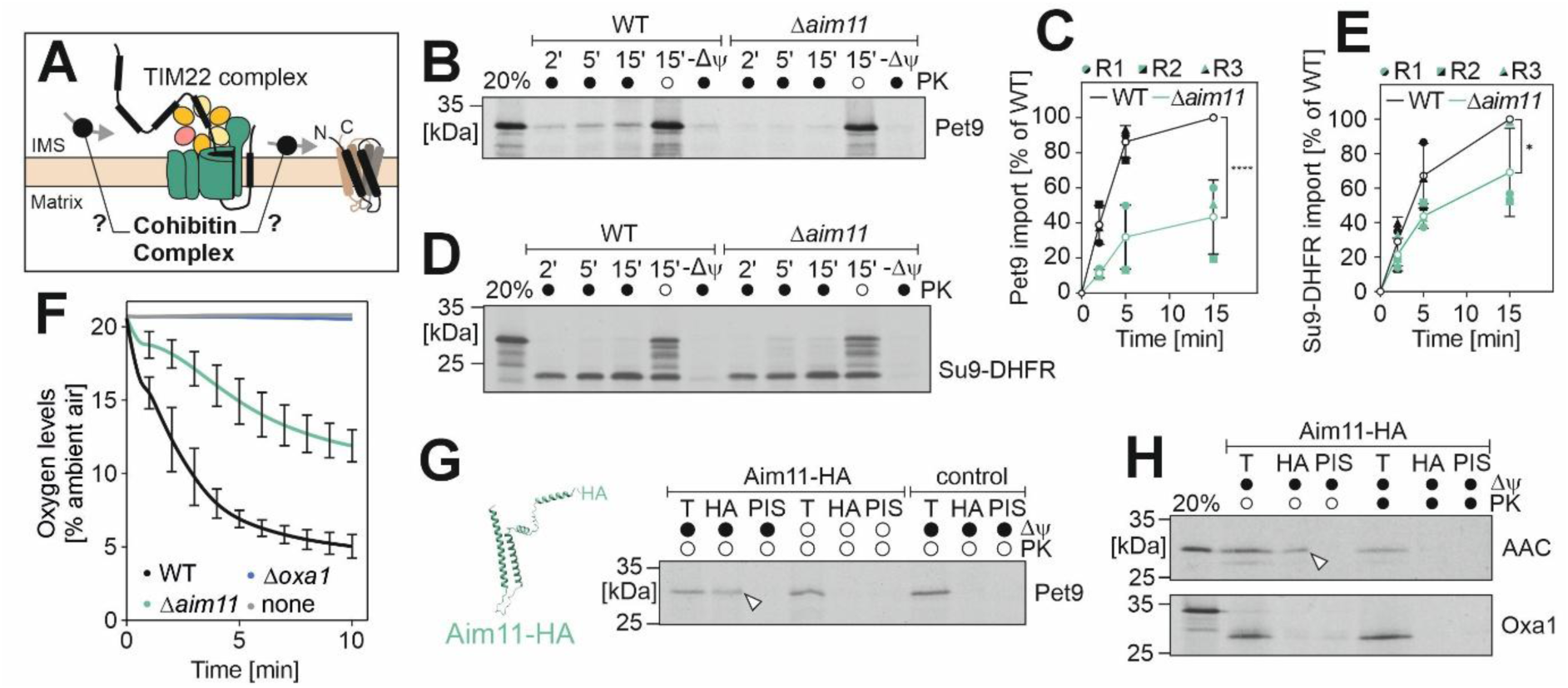
Carrier proteins interact with the cohibtin complex during membrane insertion. **(A)** Schematic representation of potential sites where the cohibitin complex acts might be upstream, downstream, or parallel to the TIM22 complex. **(B - E)** Radiolabeled Pet9 or Su9-DHFR was incubated with isolated mitochondria from wild type or *Δaim11* cells. Samples were incubated at 30°C for the indicated time points. Non-imported protein was removed by treatment with Proteinase K (PK). A control sample was treated with a mix of valinomycin, oligomycin, and antimycin A to dissipate the membrane potential (-Δψ). The 20% sample shows one-fifth of the radiolabeled protein used per time point of the import reaction. Protein levels were analyzed by SDS-PAGE. Mean values and standard deviations of three biological (independent) replicates were quantified. Statistical analysis was performed using a two-way ANOVA with Sidak’s multiple comparison test. Statistical significance was assigned as follows: p-value <0.05 = *; < 0.0001 = ****. **(F)** Oxygen consumption measurements of isolated mitochondria from the indicated strains. For wild type and *Δaim11*, the mean values and standard deviations of three biological (independent) replicates are depicted. **(G and H)** Radiolabeled Pet9, AAC, or Oxa1 were incubated with isolated mitochondria expressing Aim11-HA or empty vector as control. Samples were incubated at 30°C for 15 min with or without proteinase K (PK) addition. A control sample was treated with a mix of valinomycin, oligomycin, and antimycin A to dissipate the membrane potential (Δψ). Mitochondria were then lysed, and the extract was loaded either directly (T, equivalent to 10% total) or immunoprecipitated with anti-HA antibody-coupled magnetic beads or preimmune serum (PIS). Arrowheads depict the immunoprecipitated Pet9 or AAC signal. The 20% sample shows one-fifth of the radiolabeled protein used per sample of the import reaction.

Next, we tested whether translocation intermediates of carrier proteins are in direct contact with Aim11. We therefore employed again the strain in which Aim11 carries a C-terminal HA tag. We isolated mitochondria from this strain as well as from wild type cells for control and incubated them with radiolabeled Pet9 for 15 min. Then, the mitochondria were lysed with Triton X-100 and incubated with magnetic beads carrying anti-HA antibodies (Fig. 4G). Pet9 was co-purified with Aim11-HA (indicated by arrowhead). However, when the mitochondrial membrane potential was dissipated during the incubation to prevent Tim22-mediated insertion of carrier proteins, no radiolabeled Pet9 was co-isolated with Aim11-HA. The same pattern was seen upon incubation with the ATP/ADP carrier of *Neurospora crassa*, a commonly used model protein in the field (AAC, Fig. 4H). Thus, Aim11 apparently forms a transient complex with carrier proteins during or after their membrane potential-dependent insertion into the inner membrane.

### Aim11 promotes the TIM22-mediated carrier insertion into the inner membrane

The reduced carrier import rates in *Δaim11* mitochondria might arise from impaired translocation through the TOM complex or defects in the TIM22-mediated insertion into the inner membrane. Since carrier proteins are not made with presequences that are cleaved by the matrix processing peptidase, the *in vitro* import assays do not directly reveal the stage at which the import process is affected. We therefore generated a variant of Pet9 into which we engineered a cleavage site for the hepatitis C virus protease (HCVP) into the N-terminal matrix loop (Pet9 with cleavage site, Pet9^CS^) and expressed the HCVP enzyme with a matrix-targeting sequence in wild type and *Δaim11* cells (mt-HCVP, Fig. 5A). Endogenous yeast proteins do not carry HCVP cleavage sites and the expression of this protease in the cytosol or the mitochondria of yeast cells has no negative consequences (Flohr et al., 2025). We isolated mitochondria from these cells and incubated them with radiolabeled Pet9^CS^ (Fig. 5B, S5). In wild type mitochondria, Pet9^CS^ was efficiently cleaved by the mitochondrial mt-HCVP but not in *Δaim11* mitochondria. In the absence of Aim11, Pet9^CS^ still reached a protease-inaccessible location, presumably representing a translocation intermediate trapped in the IMS (so-called stage 3 intermediate). This indicates that Aim11 promotes the efficient insertion of Pet9^CS^ into the inner membrane.

**Figure 5.**
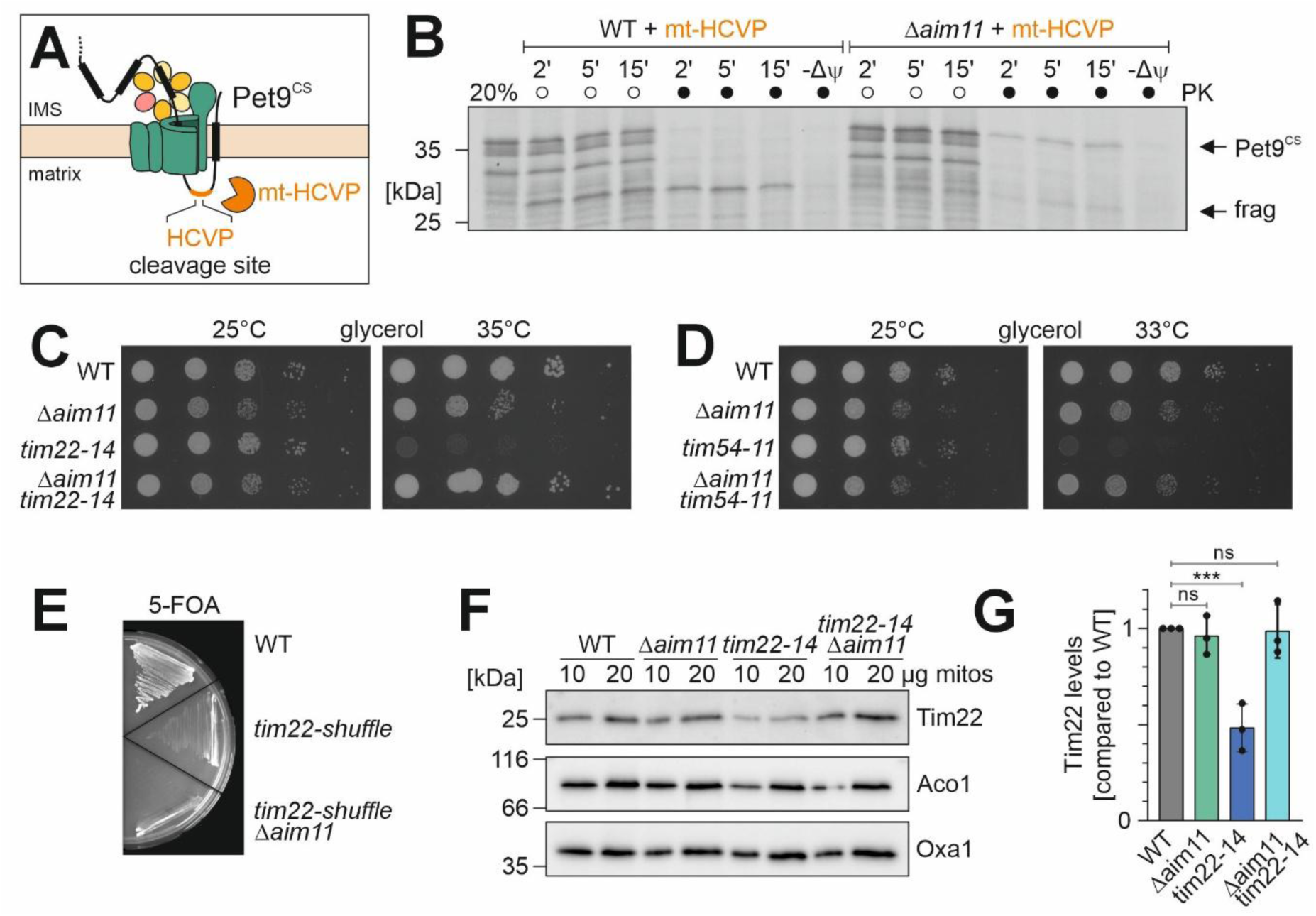
Aim11 facilitates the TIM22-dependent insertion of carriers into the inner membrane. **(A)** Schematic representation of the carrier cleavage assay. Radiolabeled Pet9 with a cleavage site (Pet9^CS^) specific to the hepatitis C virus protease (HCVP) in the first matrix-oriented loop of the imported protein is incubated with isolated mitochondria expressing the matrix-targeted protease (mt-HCVP). Successful integration into the inner membrane results in the cleavage of the protein, yielding a characteristic fragment. **(B)** Radiolabeled Pet9^CS^ was incubated at 30°C for the indicated time points with isolated mitochondria from wild type and Aim11 deletion strains expressing matrix-targeted protease. Non-imported protein was removed by treatment with Proteinase K (PK). One sample was treated with a mix of valinomycin, oligomycin, and antimycin A to dissipate the membrane potential (-Δψ). The 20% sample shows one-fifth of the radiolabeled protein used per time point of the import reaction. **(C and D)** The indicated strains were grown in galactose medium at 25°C to mid-logarithmic phase before tenfold serial dilutions were dropped onto the indicated plates and incubated at 25°C, and 33°C or 35°C. **(E)** Deletion of *AIM11* was introduced in a *Δtim22* strain using a plasmid shuffling strategy. Growth on plates containing 5-fluoroorotic acid (5-FOA) was evaluated by streaking the indicated strains. **(F and G)** Cells from the indicated strains were grown in galactose medium at 35°C, and crude mitochondria were isolated. Prior to SDS-PAGE, the mitochondrial extract was heated to 35°C for 10 min. Tim22 levels were visualized by Western blotting. Aco1 and Oxa1 served as loading controls. Mean values and standard deviations of three biological (independent) replicates were quantified. Statistical analysis was performed using a two-way ANOVA with Sidak’s multiple comparison test. Statistical significance was assigned as follows: p-value < 0.001 = ***.

Since the TIM22 complex serves as an insertase for carrier proteins (Horten et al., 2020; Kizmaz and Herrmann, 2023; Kizmaz et al., 2024; Koehler et al., 1998; Sirrenberg et al., 1996) we tested the genetic interaction of *AIM11* with components of the TIM22 complex. To this end, we deleted *AIM11* in yeast strains carrying temperature-sensitive alleles of the essential TIM22 subunits Tim22 and Tim54 (Fig. 5C, D)(Wagner et al., 2008). Surprisingly, the loss of Aim11 in these mutants largely suppressed their growth defect at restrictive temperatures. Apparently, the presence of Aim11 renders these strains unable to grow at higher temperatures. However, in *Δaim11* mutants, Tim22 was still an essential protein, as shuffle strains which harbor the *TIM22* gene on a *URA3* plasmid were unable to grow on 5-fluorotic acid (5-FOA) even if *AIM11* was deleted (Fig. 5E).

The Tim22 protein showed a reduced stability in the temperature-sensitive *tim22-14* mutant; however, in the absence of Aim11, Tim22 was again stable in the temperature-sensitive mutant suggesting that Aim11 might promote the degradation of the functionally compromised Tim22 protein (Fig. 5F, G). Deletion of Aim11 apparently suppressed the degradation of the metastable Tim22 protein, suggesting that Aim11 directly or indirectly facilitates the degradation of inner membrane proteins.

### The cohibitin complex plays a role in quality control

Our observations that the presence of Aim11 made the insertion of carriers more efficient but can lead to the degradation of compromised TIM22 complexes inspired us to test whether Aim11 plays a role in quality control. First, we challenged the carrier import route by overexpression of an N-terminally HA-tagged Pet9 protein from a *GAL* promoter (Fig. 6A). While wild type cells tolerated the overexpression of HA-Pet9 well, the HA-Pet9 overexpression in *Δaim11* cells was highly toxic.

**Figure 6.**
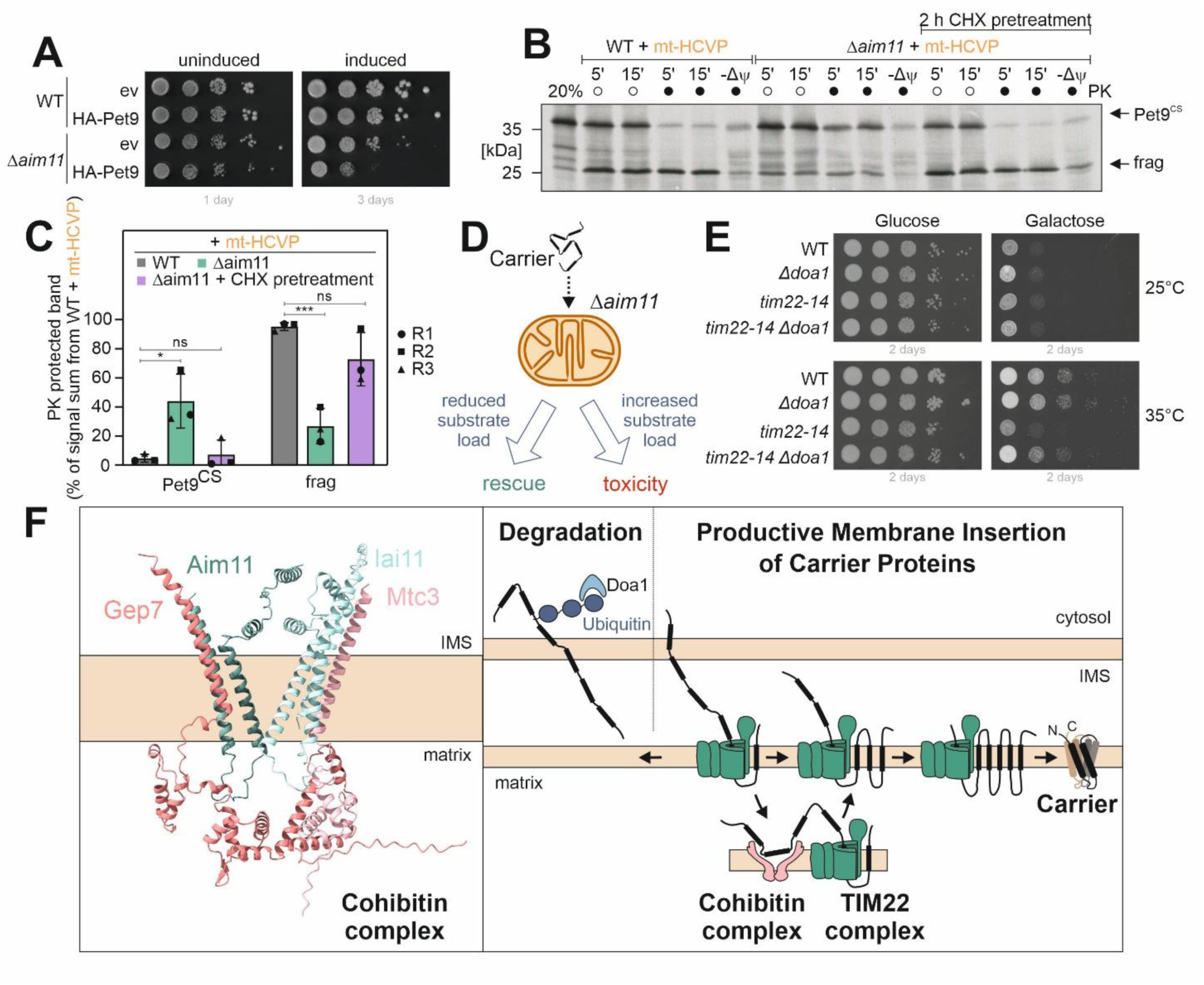
Aim11 plays a quality control function during carrier biogenesis. **(A)** The indicated strains were grown in raffinose medium at 30°C to mid-logarithmic phase before tenfold serial dilutions were dropped onto plates containing glucose (uninduced) or galactose (induced). **(B and C)** Mitochondria were isolated from wild type and Aim11 deletion strains expressing matrix-targeted hepatitis C virus protease (mt-HCVP), which were left untreated or were pretreated for 2 h with 50 µg/ml cycloheximide. Radiolabeled Pet9^CS^ was incubated for the indicated time points with crude mitochondria. Non-imported protein was removed by treatment with Proteinase K (PK). One sample was treated with a mix of valinomycin, oligomycin, and antimycin A to dissipate the membrane potential (-Δψ). The 20% sample shows one-fifth of the radiolabeled protein used per time point of the import reaction. Import signals from three biological (independent) replicates were quantified. Shown are mean values and standard deviations. Statistical analysis was performed using a two-way ANOVA with Sidak’s multiple comparison test. Statistical significance was assigned as follows: p-value < 0.05 = *; < 0.001 = ***. ns, not significant. **(D)** Aim11 facilitates carrier biogenesis; schematic model summarizing our observations. **(E)** The indicated strains were grown in galactose medium at 25°C to mid-logarithmic phase before tenfold serial dilutions were dropped onto glucose or galactose plates and incubated for the indicated amount of days at 25°C and 35°C. **(F)** Schematic model of import factors and the proteasome system which regulate carrier biogenesis. The cohibitin complex is not part of the TIM22 complex but promotes the biogenesis of carriers by transiently interacting with newly synthesized substrates, resulting in productive membrane insertion by Tim22.

We speculated that Aim11 might remove stalled or non-productive translocation intermediates from the TIM22 complex and therefore tested whether inhibition of protein synthesis in the *Δaim11* cells for 2 h prior to the isolation of mitochondria would mitigate their insertion defect. Indeed, we observed that the abstention from newly imported proteins reduces the importance of Aim11 (Fig. 6B, C), suggesting that Aim11 is particularly important if the system is charged with incoming proteins. Apparently, the cohibitin complex is critical for the efficient import of carrier proteins and counteracts the accumulation of toxic translocation intermediates (Fig. 6D).

To find out which protease might be responsible in this context, we deleted candidate genes in the temperature-sensitive *tim22-14* mutant. Thereby we observed that the absence of Doa1 phenocopied the suppression phenotype that we had found in the *Δaim11* mutant (Fig. 6D). Doa1 is a ubiquitin-binding docking factor that recruits Cdc48 to ubiquitinated proteins on the mitochondrial surface (Wu et al., 2016). This suggests that Aim11, potentially together with the other subunits of the cohibitin complex, plays a quality control function for the TIM22-mediated insertion of carrier proteins, potentially in conjunction with components of the ubiquitin-proteasome system (Fig. 6F).

## Discussion

Carrier proteins of the SLC25 family are essential for the transport of nucleotides, metabolites and metals across the inner mitochondrial membrane. All 35 (yeast) and 53 (human) members of the family contain six transmembrane domains that have to be inserted pairwise into the inner membrane by the TIM22 translocase (Brandner et al., 2005; Endres et al., 1999; Horten et al., 2020). Here, we identified Aim11 as a so far uncharacterized inner membrane protein that supports the import and membrane insertion of carrier proteins.

Owing to their high abundance and hydrophobic nature, carriers are highly aggregation-prone. Defects in the import of carriers cause cytotoxicity in yeast and humans (Wang and Chen, 2015). Presumably as a consequence of their toxic effects on mitochondrial protein biogenesis and function, point mutations in the human adenine nucleotide translocator ANT1 cause autosomal dominant Progressive External Ophthalmoplegia (adPEO) and different forms of myopathies (Kaukonen et al., 2000; Kawamata et al., 2011; Mishra et al., 2023). In this study, we identified Aim11 and other members of the cohibitin complex as quality control factors that support the import and inner membrane insertion of carrier proteins. The deletion of Aim11 renders cells hypersensitive to carrier overexpression, potentially due to a jamming of the biogenesis pathway. A quality control function of Aim11 is also supported by our observation that the import defect of *Δaim11* mitochondria can be partially mitigated if protein synthesis is blocked for 2 h prior to the preparation of mitochondria. It is evident that the cohibitin complex increases the efficiency of carrier insertion. The AlphaFold 3 models of this complex show a membrane-embedded structure that opens the lipid bilayer of the inner membrane towards the IMS. This opening might represent a binding site for partially imported carrier proteins (Fig. 6F), which is in line with the observed direct interaction of Aim11 with carrier proteins during their import but not at steady state.

While we worked on this manuscript, a preprint was published that also reported about Aim11 (Pedroza-Dávila et al., 2025). The authors also identified the four constituents of the cohibitin complex and proposed a role in the assembly of cytochrome oxidase. Interestingly, the authors also identified TMEM242 as a human homolog of Aim11, suggesting that the cohibitin complex is evolutionary conserved.

Whereas the deletion of Aim11 and other cohibitin subunits only resulted in a rather mild phenotype, the cohibitin complex became almost essential upon deletion of prohibitins. These synthetic interactions point to an overlap in the function of both complexes. Many previous studies in yeast and human cells focused on the role and relevance of the prohibitin complex, also due to its clinical relevance in the context of diseases, but its precise function is not entirely clear (Tatsuta and Langer, 2017). Prohibitins form dynamic cage-like structures that compartmentalize membranes (Hong et al., 2025; Lange et al., 2025). They presumably serve as scaffolds that support the biogenesis of inner membrane proteins by facilitating the association of insertion intermediates with membrane lipids and complex partners; moreover, if the productive assembly of their clients fails, prohibitins hand them over to membrane-embedded AAA proteases (Steglich et al., 1999; Wai et al., 2016). In contrast to prohibitins, Aim11 was not found in association with inner membrane proteases. However, we observed that the deletion of Aim11 can protect compromised subunits of the TIM22 complex against degradation. Interestingly, the deletion of the Cdc48 adapter Doa1 stabilized such TIM22 mutants and phenocopied the effects observed in the absence of Aim11. This suggests that the proteasome might be involved, for example, by proteolytic removal of partially imported carrier proteins. Tim22 also uses the carrier import pathway and is inserted into the inner membrane in an TIM22-dependent manner. It seems likely that in the *tim22-54* mutant, the proteasome removes Tim22 in the cytosol before it is transported through the TOM complex. It will be exciting to study the functional interplay of Aim11 and the other cohibitin subunits with prohibitins, the TIM22 complex, and components of the ubiquitin-proteasome system in more depth in the future.

## Materials and methods

### Yeast strains and plasmids

The yeast strains used in this study are based on the wild type strain W303 (*MAT a leu2-3,112 trp1-1 can1-100 ura3-1 ade2-1 his3-11,15*) or YPH499 (*MAT a ura3-52 lys2-801 ade2-101 trp1-Δ63 his3-Δ200 leu2-Δ*), except for the genomically mNeonGreen-tagged mutants which are based on BY4741 (*MAT a his3Δ1 leu2Δ0 met15Δ0 ura3Δ0*) (Brachmann et al., 1998; Ralser et al., 2012). Yeast strains and plasmids used in this study are described in detail in the Supplemental Tables S1 and S2.

If not otherwise specified, yeast cells were grown at 30°C in yeast full medium containing 3% (w/v) yeast extract peptone (YEP) broth (Formedium LTD) and 2% of the respective carbon source (glucose (D), galactose (Gal), glycerol (G)). Strains carrying plasmids were grown in minimal synthetic medium containing 0.67% (w/v) yeast nitrogen base and 2% of the respective carbon source. For plates, 2% of agar was added to the medium.

### Tetrade dissection

Haploid cells of opposing mating types grown to mid-log phase were mixed in a 1:1 ratio, pelleted (3 min, 3000 × g), resuspended in 75 µl YPD, and plated for mating on YPD plates for 2.5 h at 30°C. Diploid cells were selected by picking approximately 30 morphologically characteristic zygotes per mating using a SporePlay+ dissection microscope (Singer Instruments). To confirm the diploid state of the resulting colonies, each colony was plated on medium containing both antibiotics, selecting for the resistance markers of the two mating partners. For the *tim22*shuffle × *Δaim11* cross, selection was performed on –URA plates supplemented with hygromycin to confirm diploid formation. Confirmed diploids were streaked onto presporulation plates (YPD containing 5% glucose) and incubated at 30°C for 24 h. Cells were then inoculated at low density into sporulation medium (1% potassium acetate, 0.005% zinc acetate) and incubated with shaking for 5–7 days at 25°C. For tetrad dissection, sporulated cells were pelleted and resuspended in 20 µl of 5 mg/ml zymolyase (R) 20T (Amsbio, 120491-1) in 1.2 M sorbitol, incubated for 10 min at 25°C, and diluted with 180 µL of 1.2 M sorbitol to digest the ascus wall. Spores were dissected using the SporePlay+ microscope and incubated for 48 h at 30 °C prior to imaging with a Vilber Fusion FX imaging system.

Resulting tetrads were analyzed for 2:2 segregation of mating type (*MAT a* or *MAT α*) and for segregation of antibiotic resistance cassettes and/or auxotrophic markers.

### Drop dilution assay

To test growth on plates, drop dilution assays were conducted. Yeast cells were grown in full or synthetic galactose media to mid-logarithmic phase. Cultures for CRISPRi samples were treated with 960 ng/ml anhydrotetracycline (ATc) for 6 h to induce *PHB1* repression. After harvesting 2 OD_600_ of cells and washing with sterile water, a 1:10 serial dilution (starting OD = 0.5) was prepared. 3 µl of the dilutions were dropped on agar plates to determine growth differences. Plates were incubated at 30°C if not otherwise specified. Pictures of the plates were taken after 2 to 4 days of incubation.

### Preparation of whole cell lysates

For whole cell lysates, yeast strains were cultivated in liquid medium to mid-logarithmic phase. 4 OD_600_ of yeast cells were harvested and washed with sterile water. Pellets were resuspended in 50 µl/OD_600_ Laemmli buffer containing 50 mM DTT. Cells were lysed using a FastPrep-24 5 G homogenizer (MP Biomedicals) with 3 cycles of 20 s, speed 6.0 m/s, 120 s breaks, glass beads (Ø 0.5 mm) at 4°C. Lysates were heated at 96°C for 5 min and stored at −20°C.

### Antibodies

Antibodies against Sod1, Oxa1, Cmc1, Mrpl36 and Rpl6b were raised in rabbits using recombinant purified proteins. The anti-rabbit secondary antibody was obtained from Bio-Rad (Goat Anti-Rabbit IgG (H+L)-HRP Conjugate #172-1019). The antibodies against Aco1, Tim22 and Atp1 were raised in rabbits and were gifts from the Ophry Pines lab, the Nils Wiedemann lab and Marie-France Giraud from CNRS in Bordeaux, respectively. The horseradish peroxidase-coupled HA, FLAG and STRP antibodies were ordered from Roche (Anti-HA-Peroxidase, High Affinity (3F10)), Sigma Aldrich (#F3165), and Abcam (ab7403), respectively. Antibodies were diluted in 5% (w/v) nonfat dry milk in 1x TBS buffer (10 mM Tris/HCl pH 7.5, 150 mM NaCl).

### RNA isolation and real-time quantitative polymerase chain reaction (qRT-PCR)

For total RNA extraction, yeast strains were cultivated in synthetic media to exponential growth phase. 4 OD_600_ of cells were harvested, and the RNA was extracted using the RNeasy Mini Kit (Qiagen) in conjunction with the RNase-Free DNase Set (Qiagen) according to the manufacturer’s instructions. RNA quantification was performed in a two-step RT-qPCR. First, 500 ng of extracted RNA were reverse transcribed into cDNA using the qScript™ cDNA Synthesis Kit (Quantabio). Then, to measure relative mRNA levels, the iTaq Universal SYBR Green Supermix (Bio-Rad) was used with 2 µl of a 1:10 dilution of the cDNA sample. Measurements were performed in technical triplicates with the CFX96 Touch Real-Time PCR Detection System (Bio-Rad). Calculations of the relative mRNA expressions were conducted following the 2-ΔΔCt method (Livak and Schmittgen, 2001). Due to its stability, the housekeeping gene *ACT1* was used for normalization.

### Isolation of mitochondria

Mitochondria were isolated as initially described by Daum et al. (Daum et al., 1982). Yeast cells were propagated in rich or selective galactose or lactate media at 30°C to exponential phase. The temperature-sensitive *tim10-2* and *tim22-14* strains were grown at 25°C and 33°C, respectively, to allow sufficient replication while exhibiting their corresponding phenotypes in subsequent import experiments. Cells resulting from a 2 l cultivation were harvested (5 min, 2,000 × g, 25°C), washed with deionized water, and treated with 2 ml per g wet weight MP1 buffer (10 mM Tris pH unadjusted, 100 mM DTT) for 10 min at 30°C. After washing with 1.2 M sorbitol, cells were resuspended in 6.7 ml per g wet weight MP2 buffer (20 mM potassium phoshpate buffer pH 7.4, 1.2 M sorbitol, 3 mg/g wet weight zymolyase 20T from Seikagaku Biobusiness) and incubated for 1 h at 30°C. Spheroplasts were collected via centrifugation at 4°C and resuspended in 13.4 ml/g wet weight ice-cold homogenization buffer (10 mM Tris/HCl pH 7.4, 1 mM EDTA pH 8, 0.2% fatty acid free bovine serum albumin, 1 mM phenylmethylsulfonyl fluoride, 0.6 M sorbitol). Spheroblasts were disrupted by 10 strokes with a cooled glass potter. Cell debris was removed by several centrifugations at 1,500 × g for 5 min. To collect mitochondria, the supernatant was centrifuged at 12,000 × g for 12 min. Mitochondria were initially resuspended in 2 ml SH buffer (0.6 M sorbitol, 20 mM HEPES/KOH pH 7.4) and, after another centrifugation at 12,000 × g for 12 min, taken up in 200-400 µl SH buffer (volume depends on the size of the pellet). The spectrophotometer DS-11 FX+ (DeNovix) was used to determine the protein concentration and purity. Mitochondria were diluted to a protein concentration of 10 mg/ml. Aliquots were snap-frozen in liquid nitrogen and stored at −80°C.

### Import of radiolabelled precursor proteins into mitochondria

To prepare radiolabelled (^35^S-methionine) proteins for import experiments, the TNT Quick Coupled Transcription/Translation Kit from Promega was used according to the instructions of the manufacturer. To determine the ability of proteins to be imported into mitochondria, *in vitro* import assays were conducted. Mitochondria were resuspended in a mixture of import buffer (500 mM sorbitol, 50 mM HEPES pH 7.4, 80 mM KCl, 10 mM Mg(OAc)_2_, 2 mM KPi) with 2 mM ATP and 2 mM NADH to energize them for 10 min at 30°C. The import reaction was started by addition of the radiolabelled lysate (1% final volume). Import was stopped after different time points by transferring the mitochondria into cold SH buffer (0.6 M sorbitol, 20 mM HEPES pH 7.4) or 20 mM HEPES pH 7.4 (swelling conditions). The remaining precursors outside of the mitochondria were removed by proteinase K (PK) treatment for 30 min. 2 mM PMSF was added to stop protein degradation. After centrifugation (15 min, 30,000 x g, 4°C), the supernatant was removed. Pellets were resuspended in SH buffer containing 150 mM KCl and centrifuged again (15 min, 30,000 x g, 4°C). Pellets were then lysed in sample buffer (2% sodium dodecyl sulfate, 10% glycerol, 50 mM dithiothreitol, 0.02% bromophenolblue, 60 mM Tris/HCl pH 6.8) and heated to 96°C for 5 min. Samples were run on a 16 % SDS-gel, blotted onto a nitrocellulose membrane and visualized with autoradiography.

### Carbonate extraction

Radiolabeled proteins were imported *in vitro* for 15 min as described above. After the first centrifugation, the supernatant was discarded. Into one aliquot, 500 μl SH/KCl buffer with 2 mM PMSF was added, two were treated with 500 μl 0.1 M Na_2_CO_3_ pH 10 and unadjusted pH with 2 mM PMSF to extract membrane proteins with mild and more harsh conditions, respectively, and one was treated with 500 μl 0.5% TX100 with 2 mM PMSF. The samples were kept on ice for 30 min and resuspended multiple times. After centrifugation (20 min, 30,000 × g, 4°C), the supernatant was transferred into a new reaction tube. The pellet was centrifuged shortly to discard remaining liquids and lysed in 25 μl sample buffer for 5 min with agitation. Proteins in the supernatant were precipitated with 100 μl 72% trichloroacetic acid for 2 h at −70°C. Samples were then centrifuged (10 min, 30,000 × g, 4°C), the pellet washed with 500 μl ice-cold acetone, and centrifuged again (10 min, 30,000 × g, 4°C). The pellet was dried for 10 min at 37°C with an open lid. Then, the pellet was lysed in 25 μl sample buffer for 5 min with agitation. Samples were run on an 18% SDS gel, blotted onto a nitrocellulose membrane, and visualized by autoradiography.

### Co-immunoprecipitation after *in vitro* import

Radiolabeled proteins were imported in vitro for 15 min as described above. After the second centrifugation, the pellet was resuspended in 300 µl lysis buffer (0.1% TX100, 150 mM NaCl, 20 mM HEPES/KOH pH 7.4, 5 mM EDTA, 1 mM PMSF). The samples were kept on ice and resuspended multiple times. After centrifugation (15 min, 30,000 × g, 4°C), 12.5 µl of the supernatant was resuspended in the same volume of 2 × sample buffer (total sample) and the rest split into two aliquots of 125 µl each. 2 µl HA or preimmune serum were pre-coupled to 0.25 mg equilibrated magnetic protein A beads (Pierce™ protein A magnetic beads #88846) in 175 µl lysis buffer for 30 min at 4°C. The samples were incubated with the antibody-bound beads in lysis buffer, tumbling end-over-end for 1 h at 4°C. The supernatant was discarded, and the beads were washed once with 500 µl ice-cold lysis buffer and twice with 500 µl 20 mM Tris/HCl pH 7.4. Immunoprecipitated samples were taken up in 25 µl sample buffer and boiled for 5 min at 96°C. Samples were run on a 16% SDS gel, blotted onto a nitrocellulose membrane, and visualized by autoradiography.

### Fluorescence microscopy and image analysis

Yeast cells were grown in galactose media to mid-log phase, and 1 OD_600_ was harvested by centrifugation (1 min, 16,000 × g). Cell pellets were resuspended in 1 ml staining solution (5 nM MitoTracker™ Orange CMTMRos (Invitrogen®) in PBS (136.7 mM NaCl, 2.7 mM KCl, 10 mM Na2HPO4, 1.8 mM KH2PO4 adjusted to pH 7.4)) for 5 min at RT. To avoid photobleaching, cells were shielded from the light where possible. Cells were again pelleted, washed 1× with PBS, and taken up in 30 µl PBS. 3 µl of the cell suspension was pipetted onto a glass slide and covered with a cover slip. Manual microscopy was performed using a Leica Dmi8 Thunder Imager. Images were acquired using an HC PL APO 100x/1,44 Oil UV objective with Immersion Oil Type A 518 F in 16-bit format. Excitation was performed at 510 nm for mNeonGreen with an emission filter at 535 nm and 555 nm for MitoTracker Orange with an emission filter at 590 nm. The final pixel size was 132.09 µm × 132.09 µm.

Image analysis was done with the LAS X software, and further processing of images was performed in Fiji/ImageJ as follows: A background subtraction with the rolling ball set to 25 pixels was performed. Brightness and contrast were set to the same settings between samples, except for the MitoTracker Orange signal, since staining efficiency varied between samples. For spatial cross-correlation analysis, fluorescent intensity profiles were measured. Therefore, a line was drawn across the object to be analyzed (in this case, across mNG and MitoTracker orange signals). Profiles were exported as .csv files and min-max normalized.

### Oxygen consumption measurements

Oxygen consumption measurements of isolated mitochondria were conducted using the FirePlate-O2 oximeter by PyroScience. Therefore, 50 µg isolated mitochondria were taken up in 95 µl room temperature SH buffer (0.6 M sorbitol, 20 mM HEPES pH 7.4) per well in an oxygen sensor microplate. Samples were mixed with 2 mM NADH in 100 µl SH buffer using the RAININ pipetting robot (SelectScience), and optical measurements of oxygen levels were immediately started by placing the oxygen sensor microplate with a dark cover lid on the FirePlate-O2 reader. For each sample, the oxygen levels for technical triplicates were measured for 10 min.

### Sample preparation and mass spectrometric (MS) identification of proteins

For desthiobiotin labeling and affinity purification, yeast cells were inoculated in quadruplicates (n = 4) in selective lactate medium with low biotin levels (51.2 pM biotin). Cells were diluted in SLac medium without biotin and starved for 15 h. Expression of the constructs was induced by the addition of 0.5% galactose for 1 h, followed by the addition of 100 µM biotin or desthiobiotin for 3 h to start the labeling reaction. Cells equal to 10 OD600 were harvested by centrifugation (5,000 × g, 5 min), washed with ice-cold ddH2O and resuspended in 200 µl lysis buffer (2% SDS in 1 × TBS, 150 mM NaCl, 0.5 mM EDTA, 1 × EDTA-free protease inhibitor mix (cOmplete mini protease inhibitor cocktail, #04693159001, Roche). Cell lysis was performed at 4°C using a FastPrep™-24 5G homogenizer with 3 cycles of 20 s and 8.0 m/s speed, and 120 s breaks in between. Following another centrifugation, lysates were transferred to pre-cooled reaction tubes and stored at −70°C. To remove free biotin and desthiobiotin, proteins were precipitated with 1.2 ml ice-cold acetone at −20°C for 2 h. Samples were then centrifuged (10 min, 13,000 × g, 4°C) and the supernatant discarded. The sample was resolved in 900 µl ice-cold acetone and 100 µl 1 × TBS, incubated at −20°C for 2 h, and centrifuged again (10 min, 13,000 × g, 4°C). The pellet was dried for 5 min on an ice block with an open lid and dissolved in 300 µl resolubilization buffer (8 M urea, 50 mM ammonium bicarbonate, 1 mM PMSF, 1 × EDTA-free protease inhibitor mix). 0.5 mg magnetic STRP beads (Pierce™ streptavidin magnetic beads #88816) were equilibrated in 2 M urea in 1 × TBS and 1 × EDTA-free protease inhibitor mix and added to the lysed samples, tumbling end-over-end for 1 h at RT. The beads were washed 3 × with wash buffer I (150 mM NaCl, 50 mM Tris/HCl pH 7.5,5% glycerol, 0.05% NP40) and 2 × with wash buffer II (150 mM NaCl, 50 mM Tris/HCl pH 7.5, 5% glycerol). Samples were then digested with 5 ng/µl sequencing grade modified trypsin (Promega, #V5111) for 1 h at RT in 50 µl elution buffer I (2 M urea, 50 mM Tris/HCl pH 7.5, 1 mM DTT), before adding fresh trypsin for a final concentration of 15 ng/µl for 10 min. Finally, 50 µl elution buffer II (2 M urea, 50 mM Tris/HCl pH 7.5, 5 mM chloroacetamide) was added and incubated overnight in the dark at RT. Peptides were acidified to pH < 2 with trifluoroacetic acid and desalted on self-constructed StageTips containing three layers of reversed-phase Empore™ C18 disks (Rappsilber et al., 2007). C18 StageTips were activated with 100 µl methanol and equilibrated with 100 µl buffer B (0.1% formic acid, 80% acetonitrile) and 100 µl buffer A (0.1% formic acid). The acidified peptides were added onto the StageTips and washed with 100 µl buffer A. Peptides were eluted with 60 µl buffer B and dried down in a SpeedVac. Samples were resolubilized in 9 µl buffer A++ (0.1 % formic acid, 0.01% trifluoracetic acid in MS-grade water).

For co-immunoprecipitation on HA beads, yeast cells were grown in triplicates (n = 3) in selective galactose medium, and crude mitochondria were isolated as described before. 500 µg isolated mitochondria were taken up in 1 ml ice-cold lysis buffer (10 mM Tris/HCl pH 7.4, 150 mM NaCl, 0.5 mM EDTA, 1% (w/v) digitonin, 1 mM PMSF, 1 × EDTA-free protease inhibitor mix). Cell lysis was performed at 4°C using an analogue Genie® Disruptor (Scientific Industries) for 10 min. Samples were then centrifuged (1 min, 2,000 × g, 4°C). 3 µl HA antibody (Sigma Aldrich, #SAB2702249) was pre-coupled to 0.4 mg equilibrated magnetic protein A beads (Pierce™ protein A magnetic beads #88846) in lysis buffer for 30 min at 4°C. Lysed samples were incubated with the equilibrated beads by end-over-end rotation for 1 h at 4°C. The beads were washed 2 × with wash buffer I (150 mM NaCl, 50 mM Tris/HCl pH 7.5, 5% glycerol, 0.1% (w/v) digitonin) and 2 × with wash buffer II (150 mM NaCl, 50 mM Tris/HCl pH 7.5, 5% glycerol). For on-bead digestion, elution buffer I (2 M urea, 50 mM Tris/HCl pH 7.5, 1 mM DTT, and 5 ng/µl trypsin) was added, and samples were incubated for 1 h at RT. The supernatant was transferred into a fresh reaction tube. For a second elution step, elution buffer II (2 M urea, 50 mM Tris/HCl pH 7.5, 5 mM chloroacetaldehyde, 5 ng/µl trypsin) was added to the beads, and the samples were incubated for 30 min at RT. The supernatant of the first and second elution steps were combined and incubated overnight in the dark at 37°C. Peptides were acidified and cleaned up with StageTips containing three layers of Empore™ C18 disks.

For quantitative whole proteome comparisons of strains carrying CRISPRi plasmids, cells were harvested in quadruplicates (n = 4) after 24 h of induction with 960 ng/ml ATc. 10 OD_600_ cells were harvested by centrifugation (5,000 × g, 5 min, RT), washed twice with ice-cold MS-grade H2O and transferred into pre-cooled reaction tubes. After another centrifugation (16,000 × g, 1 min, 4°C), the supernatant was removed, the pellets snap-frozen in liquid nitrogen and stored at −70°C. Cell pellets were resuspended in 200 µl lysis buffer (6 M guanidinium chloride, 10 mM TCEP-HCl, 40 mM chloroacetamide, 100 mM Tris/HCl pH 8.5 in MS-grade water). Cell lysis was performed at 4°C using an analogue Genie® Disruptor (Scientific Industries) for 15 min, followed by two centrifugations at 16,000 × g for 5 min and 4°C. In between, the supernatant was transferred to fresh reaction tubes to remove any remaining glass beads. Protein concentrations were measured using the Pierce BCA Protein Assay Kit (Thermo Scientific, #23225). For protein digestion, 25 µg of protein was diluted 1:10 in LT-digestion buffer (10% acetonitrile, 25 mM Tris/HCl pH 8.8) and 1:50 (w/w) trypsin and lysyl endopeptidase R (Wako, #125-05061) added. Samples were incubated overnight at 37°C and 700 rpm. Peptides were acidified to pH < 2 with trifluoroacetic acid and desalted on self-constructed StageTips containing three layers of reversed-phase Empore™ SDB-RPS disks (Cat #2241)(Rappsilber et al., 2007). SDB-RPS StageTips were activated with 100 µl acetonitrile and equilibrated first with 100 µl of 1% trifluoracetic acid in 30% methanol and second with 100 µl of 0.2% trifluoracetic acid. The acidified peptides were added onto the StageTips and washed twice with 100 µl of 0.2% trifluoracetic acid. Peptides were eluted with 60 µl of 5% ammonia in 80% acetonitrile and dried down in a SpeedVac. Samples were resolubilized in 12 µl buffer A++ (0.1 % formic acid, 0.01% trifluoracetic acid in MS-grade water).

Peptides were separated on 50 cm columns with a 75 µm inner diameter that were packed in-house with C18 resin (ReproSil-Pur 120 C18 AQ, 1.9 µm; Dr. Maisch) using an EASY nLC 1200 system (Thermo Fisher Scientific, Germany). The eluted peptides were directly sprayed into a Q Exactive HF mass spectrometer (Thermo Fisher Scientific, Germany). Separation was achieved using a 3-hour linear gradient from 2% to 95% buffer B*, consisting of buffer A (0.1% formic acid) and buffer B (80% acetonitrile, 0.1% formic acid). Mass spectrometric data were acquired in data-dependent acquisition mode and processed with MaxQuant software version 2.6.4.0 (Cox and Mann, 2008). All additional data acquisition and processing parameters are available in the datasets deposited in PRIDE (see below).

### Complexome profiling analysis

Wild type, *rho^0^* or *GAL*-Rpo41 cells were grown in galactose-containing media. 48 h before the cells were harvested, the *GAL*-Rpo41 cells were split into three cultures. One was continuously grown on galactose, one shifted to raffinose for 48 h and one to raffinose for 43 h before galactose was added for 5 h. From all cultures, cells were harvested, and mitochondria were isolated.

For complexome data analysis, protein concentrations were determined using the microBCA protein kit (Thermo Scientific). Mitochondria were solubilized with 3 g digitonin (SERVA) per gram of protein in 50 mM NaCl, 2 mM 6-aminohexanoic acid, 1 mM EDTA, and 50 mM imidazole/HCl (pH 7.0). Blue native electrophoresis was performed as described in Wittig et al. (Wittig et al., 2006). The in-gel digestions were performed essentially as previously described with slight modifications (Heide et al., 2012). Following native electrophoresis, the gel was incubated in fixing solution (50% methanol, 10% acetic acid, 100 mM ammonium acetate) for 30 min and stained with Coomassie blue. After two destaining steps with 10% acetic acid (1 h each), the gel lanes were cut into 60 even slices of about 2 mm. Every gel slice was additionally diced into smaller pieces and then transferred to a 96-well filter microplate containing 200 µl of 50% methanol, 50 mM ammonium hydrogen carbonate (pH unadjusted). The gel pieces were washed in destaining solution at room temperature under gentle agitation until complete Coomassie dye removal. Excess solution was removed by centrifugation (600 × g, 1 min at RT). In the next step, gel pieces were incubated with 120 µl of 10 mM DTT for 60 min. After removal of excess solution, 120 µl of 30 mM chloroacetamide was added to each well and removed after 45 min. Gel pieces were dried at room temperature for 45 min. The dried gel pieces were swollen in 20 µl of 5 ng/ml trypsin in 50 mM ammonium hydrogen carbonate and 1 mM CaCl_2_ (pH unadjusted) for 30 min at 4°C. Then, 150 ml of 50 mM ammonium hydrogen carbonate was added to cover the gel pieces, followed by an overnight incubation at 37°C to digest the proteins. The peptide-containing supernatants were collected by centrifugation (600 × g, 3 min, RT) into a 96-well plate. The gel pieces in the filter plate were washed once with 30% acetonitrile, 3% formic acid for 20 min to elute the remaining peptides. The combined eluates were then dried at 45°C for 3 h in a Concentrator Plus (Eppendorf). The peptides were resuspended in 20 µl of 5% acetonitrile/0.5% formic acid and stored at −20°C until mass spectrometric analysis.

Peptides from tryptic digestion were separated and analyzed by liquid chromatography–tandem mass spectrometry (LC-MS/MS) using a Q Exactive Orbitrap mass spectrometer coupled to an Easy-nLC 1000 nano-flow ultra-high-pressure liquid chromatography (UHPLC) system (Thermo Fisher Scientific), following the settings described in Guerrero-Castillo et al. (Guerrero-Castillo et al., 2017). In brief, peptides were separated on a 100 μm inner diameter × 15 cm PicoTip emitter column (New Objective) packed with ReproSil-Pur C18-AQ reverse-phase beads (3 μm particle size, 120 Å pore size; Dr. Maisch GmbH). Chromatographic separation was achieved using a linear gradient of 5%–35% acetonitrile (ACN) in 0.1% formic acid (FA) over 30 min, followed by 35%–80% ACN in 0.1% FA for 5 min at a flow rate of 300 nl/min, and a final column wash with 80% ACN for 5 min at 600 nl/min. The mass spectrometer was operated in positive ion mode, alternating automatically between full MS and data-dependent MS/MS acquisition of the top 20 most intense precursor ions (Top20 method). Full MS scans (m/z 400–1400) were acquired at a resolution of 70,000 (at m/z 200), with an automatic gain control (AGC) target of 1 × 10⁶ ions and a maximum injection time of 20 ms. MS/MS scans were acquired at a resolution of 17,500 (at m/z 200), with an AGC target of 1 × 10⁵, maximum injection time of 50 ms, and precursor isolation window of 4.0 Th. Only precursor ions with charge states z = 2 or z = 3 were selected for fragmentation by higher-energy collisional dissociation (HCD), using a normalized collision energy of 30%. A dynamic exclusion window of 60 s was applied. Internal mass calibration was performed using a lock mass ion (m/z 445.12) as described in Olsen et al. (Olsen et al., 2005).

Raw MS data from all fractions were processed using MaxQuant (v1.5.0.25). Peptide spectra were matched against the Baker’s yeast UniProt reviewed proteome (TaxID 4932, release date July 2020). The search parameters included cysteine carbamidomethylation as a fixed modification, as well as N-terminal acetylation and methionine oxidations as variable modifications. Trypsin/P was specified as the protease, allowing up to two missed cleavages. A minimum peptide length of six amino acids was set. The “match between runs” feature was enabled, with a matching time window of 2 min, and unidentified feature matching was allowed. Additionally, isoleucine was treated as equivalent to leucine. Protein abundances were determined by label-free quantification using the resulting intensity-based absolute quantification (iBAQ) values. Apparent molecular mass calibration was done using a set of known membrane and soluble proteins from yeast and bovine heart mitochondria. Data were clustered (Pearson correlation, average-linkage) and pre-processed using an in-house java-based script. Relative abundance was analyzed and visualized as heatmaps/line charts using Microsoft Excel.

### Mass spectrometric data analysis

All mass spectrometry data sets were processed using the R programming language. For affinity purification of desthiobiotinylated proteins and for co-immunoprecipitation, proteins that were identified in less than three replicates of the non-control sample (expressing the Destni-fusion protein or the HA-tagged protein respectively, n = 3) were removed, which resulted in 2691 and 127 robustly identified protein groups, respectively. For whole cell proteomics of the *Δaim11* and Phb1-depletion strains, proteins that were identified in less than four replicates per sample were removed (n = 4), which resulted in 3674 identified protein groups. Log_2_-transformed label-free quantification (LFQ) values were normalized using variance stable normalization and batch effect was removed for co-immunoprecipitation with HA beads and whole cell proteomics using the limma package. Lastly, for affinity purification of desthiobiotinylated proteins and for co-immunoprecipitation, missing values of the control samples were imputed if a protein was not measured in three replicates by sampling n = 3 values from a normal distribution (seed = 73294168). Proteins were tested for differential expression using limma for the indicated comparison of samples and p-values were adjusted for multiple testing using a Benjamini-Hochberg procedure (Benjamini and Hochberg, 1995). Test results for affinity purification of desthiobiotinylated proteins are listed in Table S3. Test results for the co-immunoprecipitation with HA-beads are listed in Table S5. Test results for the whole cell proteome of the *Δaim11* and Phb1-depletion strains are listed in Table S6. For principal component analysis, the processed LFQ intensities were used for singular value decomposition. Mitochondrial sublocalization was determined using a merged and manually curated list derived from published data. For plotting a heatmap of affected mitochondrial proteins in the *Δaim11* and Phb1-depletion strains, processed LFQ values were filtered for mitochondrial proteins and averaged per sample. Then, z-scores were calculated and centered to the wild type strain. Hierarchical clustering was performed using Canberra distances and unweighted pair group method with arithmetic mean. Results are listed in Table S7. For gene ontology analysis of desthiobiotinylated proteins, all quantified proteins were used as background set and proteins with log_2_ fold change > 0.5 and adjusted p-value < 0.05 were used as target set using the GOrilla tool (Eden et al., 2009). Results are listed in Table S4. For gene ontology analysis of the whole cell proteome of the *Δaim11* and Phb1-depletion strains, the results were filtered for the indicated comparison, and all quantified proteins were used as background set and proteins with log_2_ fold change < −0.5 and adjusted p-value < 0.05 were used as target set for the GOrilla tool. Results are listed in Table S8.

### Modelling of protein complexes

The 3D structure or protein complexes was modeled using the open-source tool AlphaFold Server, which is based on AlphaFold 3 (Abramson et al., 2024). The amino acid sequence of the proteins served as input. The resulting 3D models of the complexes were downloaded as PDB files and further edited using UCSF ChimeraX version 1.9.

## Data availability statement

All reagents and strains as well as the R scripts for the analysis of proteomics data are available upon request from the corresponding author (JMH). The mass spectrometry proteomics data (see also Tables S3, S5, S6) have been deposited to the ProteomeXchange Consortium via the PRIDE partner repository with the dataset identifier shown below.

Reviewer account details:

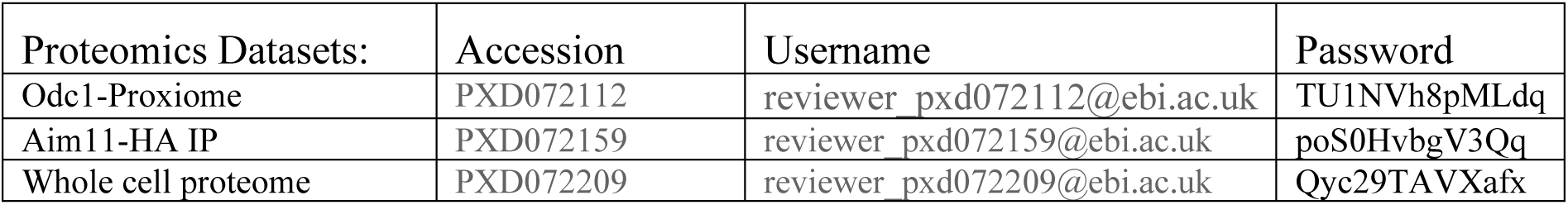

The complexome profiling dataset has been deposited in the CEDAR database under accession code **CRX51** (www3.cmbi.umcn.nl/cedar/browse/experiments/CRX51) (van Strien et al., 2021). All other MS raw data are available by the authors upon reasonable request.

## Acknowledgements

We thank Sabine Knaus for technical assistance and Ophry Pines, Nils Wiedemann and Marie-France Giraud for reagents. We thank Nils Wiedemann for the *tim22* shuffle strain and the temperature-sensitive *tim22-14* and *tim54-11* mutants. This study was financially supported by grants from the Deutsche Forschungsgemeinschaft (RTG2737 STRESSistance and SPP2453 project number 541210481 to JMH), the European Research Council (MitoCyto 101052639 to JMH) and the Landesforschungsiniative Rheinland-Pfalz BioComp.

## Abbreviations

Cohibitin: comrade-of-prohibitin
Destni: desthiobiotin ligase
DHFR: dihydrofolate reductase
HCVP: hepatitis C virus protease
IMS: intermembrane space
iMTS: internal matrix targeting signal
MTS: matrix targeting signal
TIM: translocase of the inner membrane
TOM: translocase of the outer membrane

## Conflict of Interests

The authors have no conflicts of interest.

---

**Figure S1.**
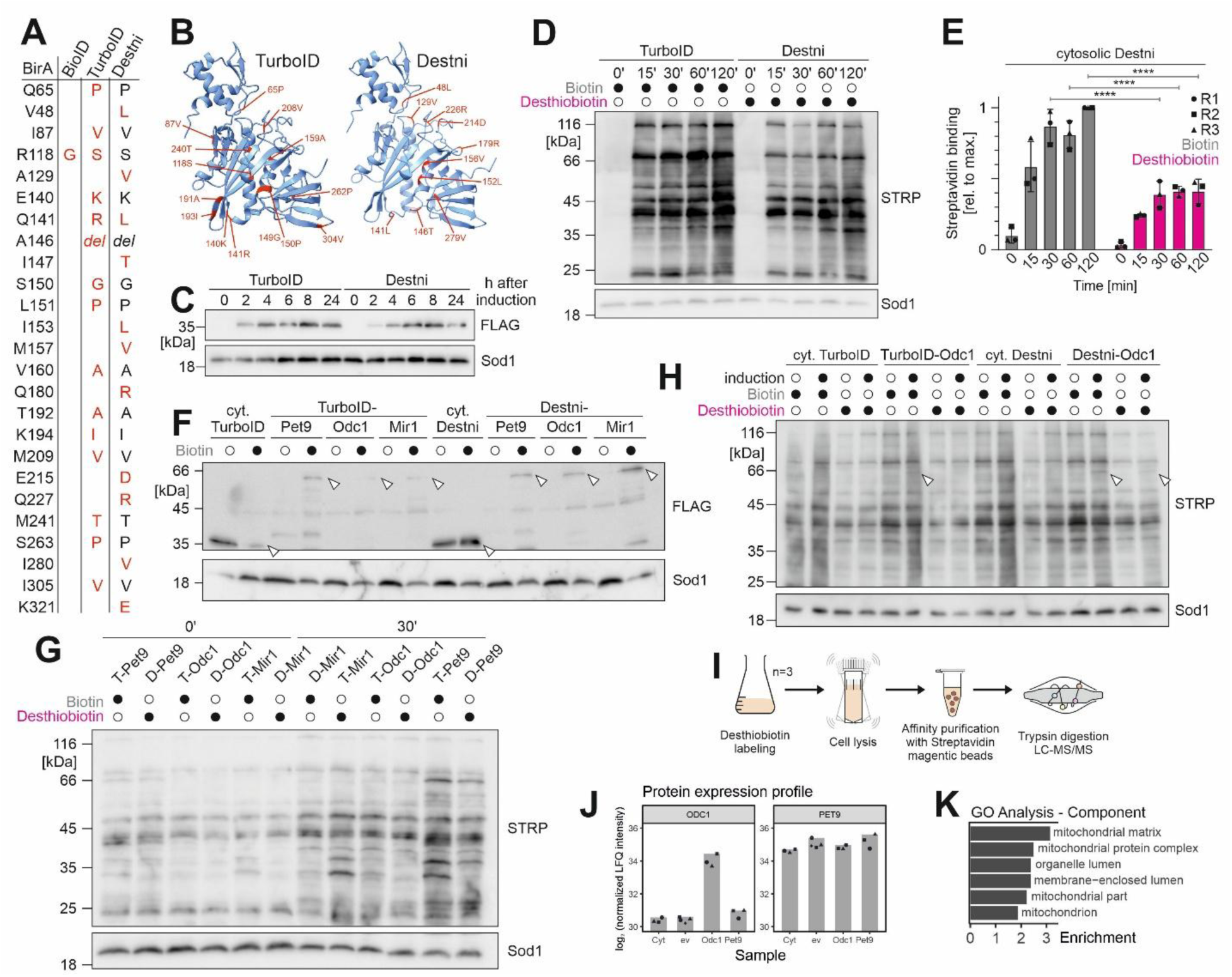
Related to Fig. 1. Destni allows to efficiently desthiobiotinylate proteins in its proximity. **(A)** Changes in the amino acid sequence in the biotin ligases BioID, TurboID and DestniD compared to *E. coli* BirA (Branon et al., 2018). All biotin ligases represent improved versions of the bacterial BirA. Residues labeled in red represent newly introduced mutations. **(B)** AlphaFold 3 prediction of TurboID and Destni. Mutated residues of TurboID relative to BioID and Destni compared to TurboID are indicated in the structures (Abramson et al., 2024). **(C)** Expression of the C-terminally FLAG-tagged cytosolic TurboID and Destni, respectively, induced by the addition of 0.5% galactose for up to 24 h. Cells were harvested and lysed at the indicated time points. Sod1 serves as a loading control. **(D and E)** Following expression of TurboID and Destni with 0.5% galactose for 4 h, respectively, the labeling reaction was started by adding 100 µM biotin or desthiobiotin for up to two hours. Cells were harvested at the indicated time points, lysed, and analyzed via Western blot. Sod1 antibody serves as a loading control. Whole lane signals from three biological (independent) replicates were quantified. Shown are mean values and standard deviations. Statistical analysis was performed using a two-way ANOVA with Sidak’s multiple comparison test. Statistical significance was assigned as follows: p-value < 0.0001 = ****. **(F)** Expression of the C-terminally FLAG-tagged cytosolic biotin ligases and fusion constructs with the carrier proteins Pet9, Odc1, and Mir1 were induced with 0.5% galactose for 4 h and protein levels were analyzed via Western blot. Arrowhead indicated the full-length constructs. Sod1 antibody serves as a loading control. **(G, H)** The indicated fusion constructs were expressed for 4 h. The labeling reaction was initiated by adding 100 µM biotin or desthiobiotin for the indicated duration. Cells were harvested, lysed, and analyzed via Western blot. Levels of biotinylated proteins were detected with streptavidin-coupled horse radish peroxidase (STRP). Sod1 served as a loading control. T, TurboID; D, Destni (**I**) Scheme for the analysis of Odc1 interactors by mass spectrometry. (**J**) Enrichment of Odc1 (left) and Pet9 (right) from the affinity purification of proximity labeling proteomics analysis. (**K**) Proteins that were explicitly enriched in the Destni-Odc1 vs. empty vector (ev) samples (log_2_ fold change > 0.5 and p-value <0.05) were analyzed by gene ontology (GO) enrichment using the GOrilla tool (http://cbl-gorilla.cs.technion.ac.il). log_2_ FC, log_2_ fold change.

**Figure S2.**
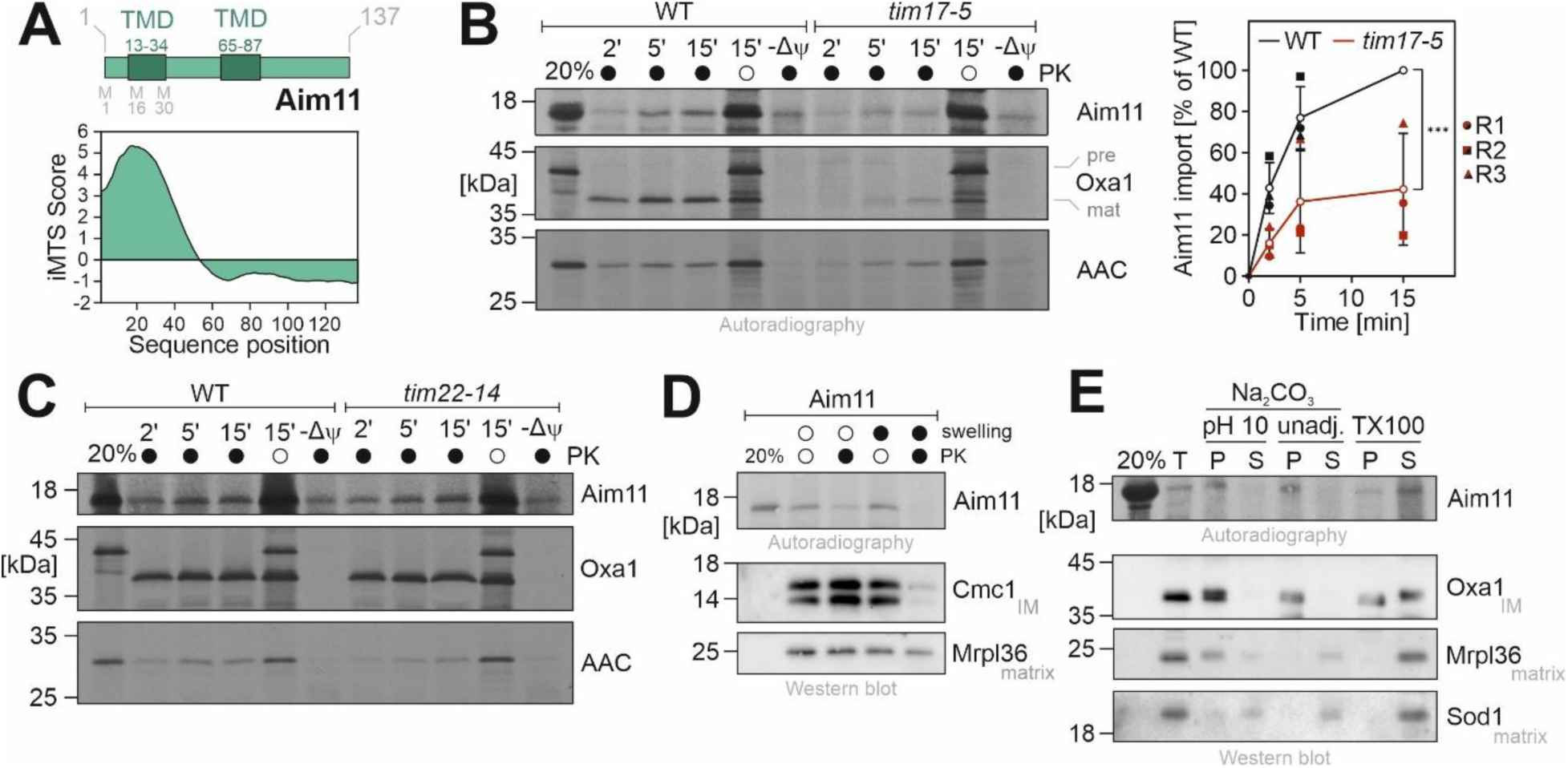
Related to Fig. 1. Aim11 is imported into the inner membrane by the TIM23 pathway. **(A)** The iMTS score was calculated for each residue in the Aim11 sequence and plotted, showing a high probability for an internal mitochondrial targeting signal around the first transmembrane domain (TMD)(Jung et al., 2024). **(B)** (Left) Aim11, Oxa1, and ATP/ADP carrier (AAC) were synthesized in reticulocyte lysate in the presence of ^35^S methionine and incubated with mitochondria isolated from wild type (WT) or *tim17-5* cells that were grown at restrictive conditions (35°C, 18 h) in lactate medium. After the times indicated, samples were treated with or without proteinase K (PK). To dissipate the membrane potential (Δψ), a mix of valinomycin, antimycin, and oligomycin was used. The 20% sample shows 20% of the radiolabeled Aim11 protein used per import reaction; pre, precursor; mat, mature. (Right) The signals of three independent Aim11 samples were used for quantification. Shown are mean values and standard deviations. Statistical analysis was performed using a two-way ANOVA with Sidak’s multiple comparison test. Statistical significance was assigned as follows: p-value < 0.001 = ***. **(C)** Aim11, Oxa1, and ATP/ADP carrier (AAC) lysates were imported into mitochondria isolated from wild type (WT) or *tim22-14* cells that were grown at restrictive conditions (35°C, 18 h) in lactate medium according to Fig. S2B. **(D)** Aim11 was imported into mitochondria isolated from wild type cells for 15 min. After import, mitochondria were incubated in iso- or hypoosmotic (swelling) buffer in the absence or presence of proteinase K (PK). Protein levels were analyzed by SDS-PAGE. Western blot signals of the inner membrane (IM) protein Cmc1 and the matrix protein Mrpl36 served as controls. **(E)** Aim11 was imported into wild type mitochondria for 15 min before mitochondria were incubated with carbonate of pH 10, of unadjusted pH (unadj.), or with 0.5% Triton X-100 (TX-100). After incubation for 30 min on ice, samples were centrifuged, and the proteins from the supernatant (S) and the pellet (P) were analyzed by SDS-PAGE. T, total. Western blot signals of the inner membrane (IM) protein Oxa1 and the matrix proteins Mrpl36 and Sod1 served as controls.

**Figure S3.**
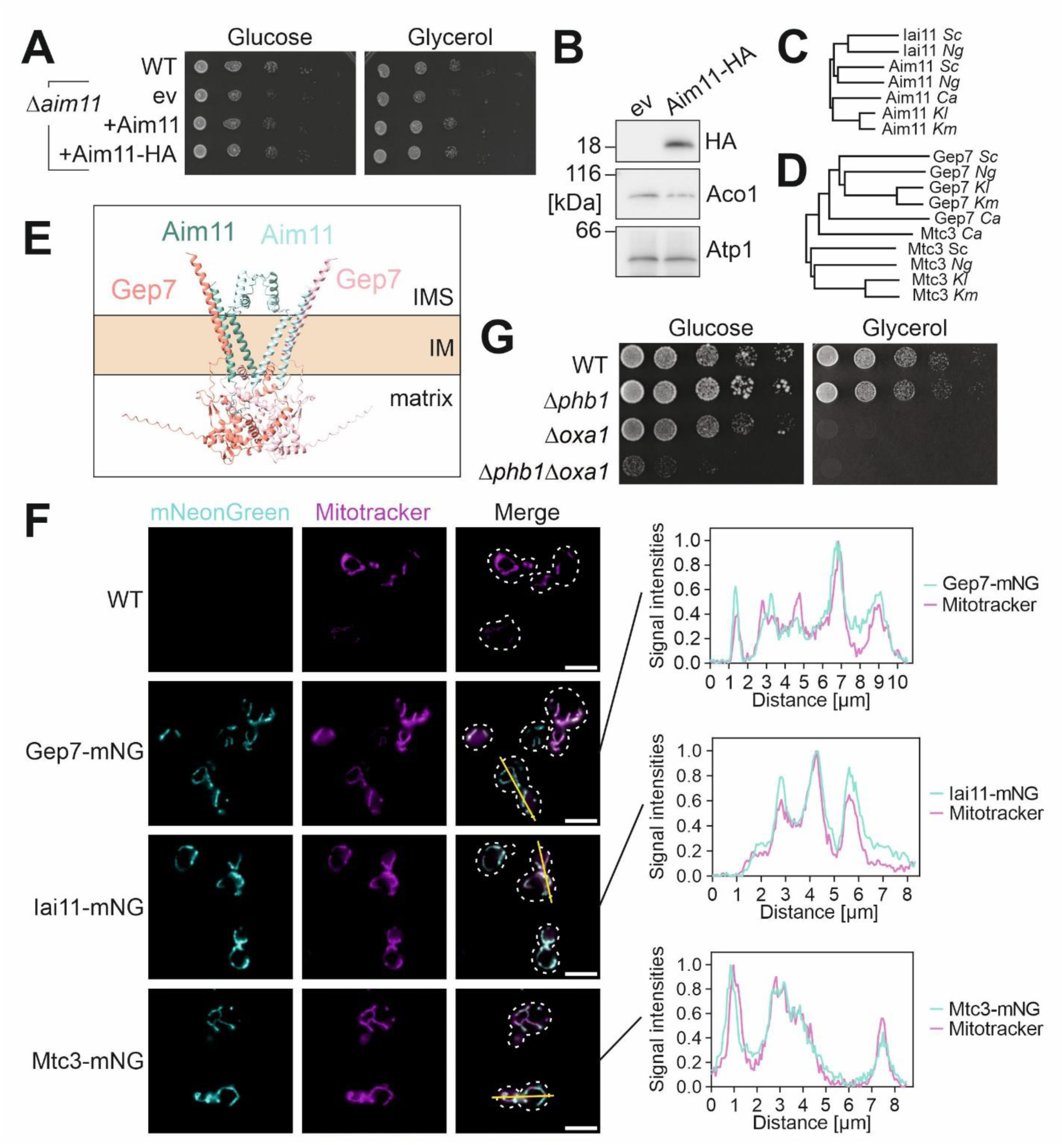
Related to Figs. 2 and 3. Gep7, Iai11 and Mtc3 are mitochondrial proteins. **(A)** Wild type (WT) and *Δaim11* cells containing plasmids for the expression of Aim11, Aim11-HA or an empty plasmid (ev) were grown to mid-logarithmic phase on galactose medium before tenfold serial dilutions were dropped on plates containing the indicated carbon sources. Plates were incubated at 30°C. **(B)** Extracts of *Δaim11* cells with Aim11-HA expression plasmid or an empty plasmid were analyzed by Western blotting. Aco1 and Atp1 served as loading controls. **(C, D)** The phylogenetic tree was calculated with Muscle multiple sequence alignment tool (Madeira et al., 2024) on the basis of the following sequences: Gep7 (*Sc*, *Saccharomyces cerevisiae* P53171; *Kl*, *Kluyveromyces lactis* QEU60090.1; *Ca*, *Candida albicans* KAF6061140.1; *Km*, *Kluyveromyces marxianus* KAG0670543.1; *Ng*, *Nakaseomyces glabratus* KAH7580541.1). Mtc3: *Sc* P53077; *Kl* XP_451055.1; *Km* KAG0679606.1; *Ng* CAG60366.1; *Ca* XP_723445.1. Aim11: *Sc* A0A815YB48; *Kl* XP_454863.1; *Ng*, *Nakaseomyces glabratus* KAJ9570390.1; *Ca* XP_710638.1; *Km* KAL2709502.1. Iai11: *Sc* P34224; *Ng* XP_444957.1. **(E)** AlphaFold 3 prediction of the (Aim11)_2_-(Gep7)_2_ complex (Abramson et al., 2024). **(F)** (Left) Wild type (WT) and genomically mNeonGreen-tagged strains were grown in full galactose medium to mid-logarithmic phase and visualized by fluorescence microscopy. Shown are the channels for mNeonGreen, Mitotracker Orange to image the mitochondrial network, and the merge. Scale bar: 5 µm. (Right) Min-max normalized plot profiles of the signals along the yellow line shown in the merge. **(G)** Wild type and the indicated deletion mutants were grown on galactose-containing media to mid-log phase before tenfold serial dilutions were dropped onto the indicated plates and incubated at 30°C.

**Figure S4.**
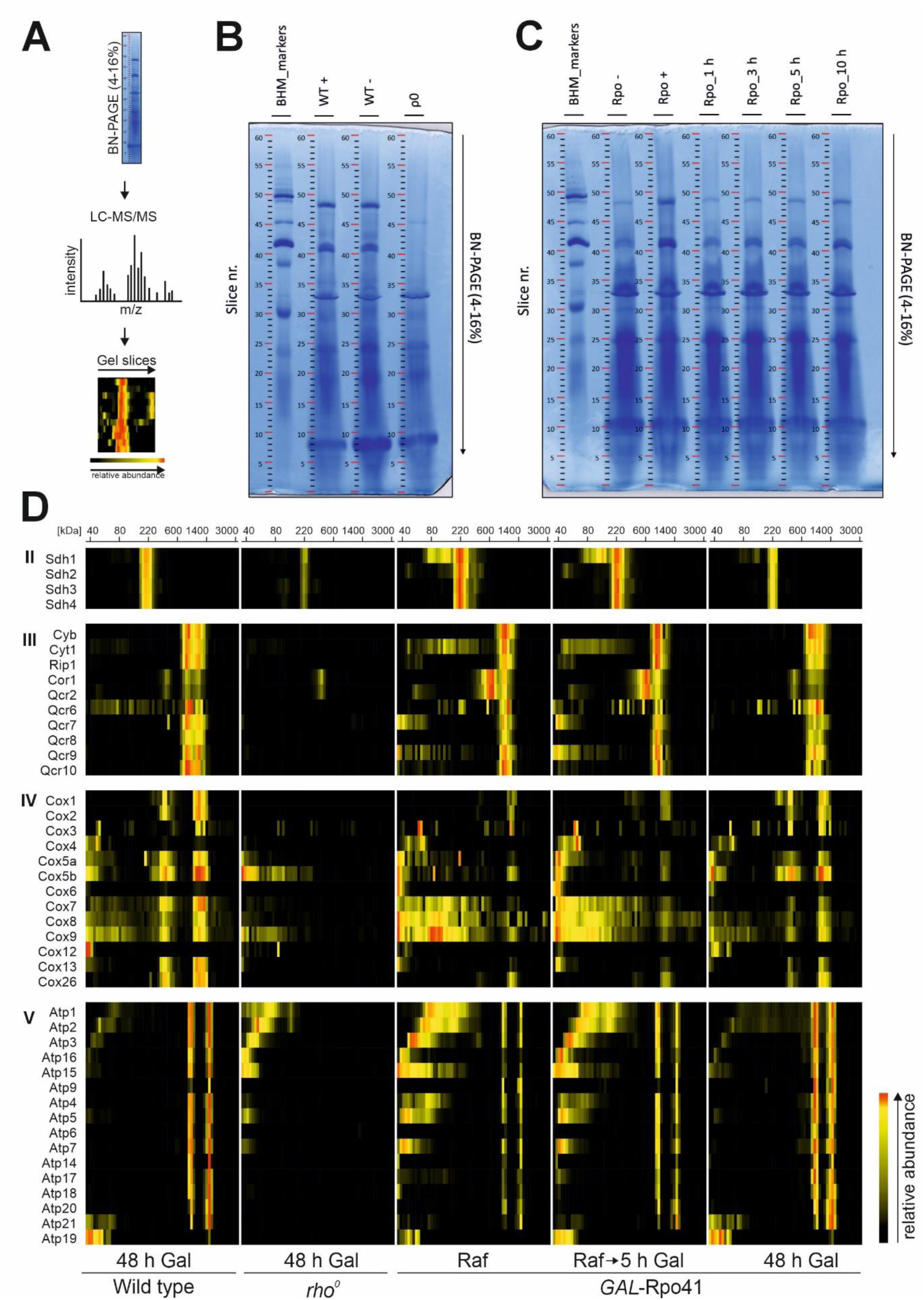
Related to Fig. 2. Complexome analysis of mitochondrial protein complexes. **(A)** Protein complexes were separated by blue-native PAGE according to their size. The gel was cut into 60 slices of even size, and each slice was processed separately by in-gel tryptic digestion followed by liquid chromatography-tandem mass spectrometry. Migration patterns were reconstructed by combining the identified relative protein abundance from each mass spectrometry analysis. **(B and C)** Blue-native PAGE of the tested strains. **(C)** Complexome profiling heatmap of OXPHOS complexes II, III, IV and V under steady state conditions. The migration profiles of components of OXPHOS complexes were assembled manually and color-coded by normalizing the relative abundance for each subunit separately taking the maximal value across the experimental condition as 1.

**Figure S5.**
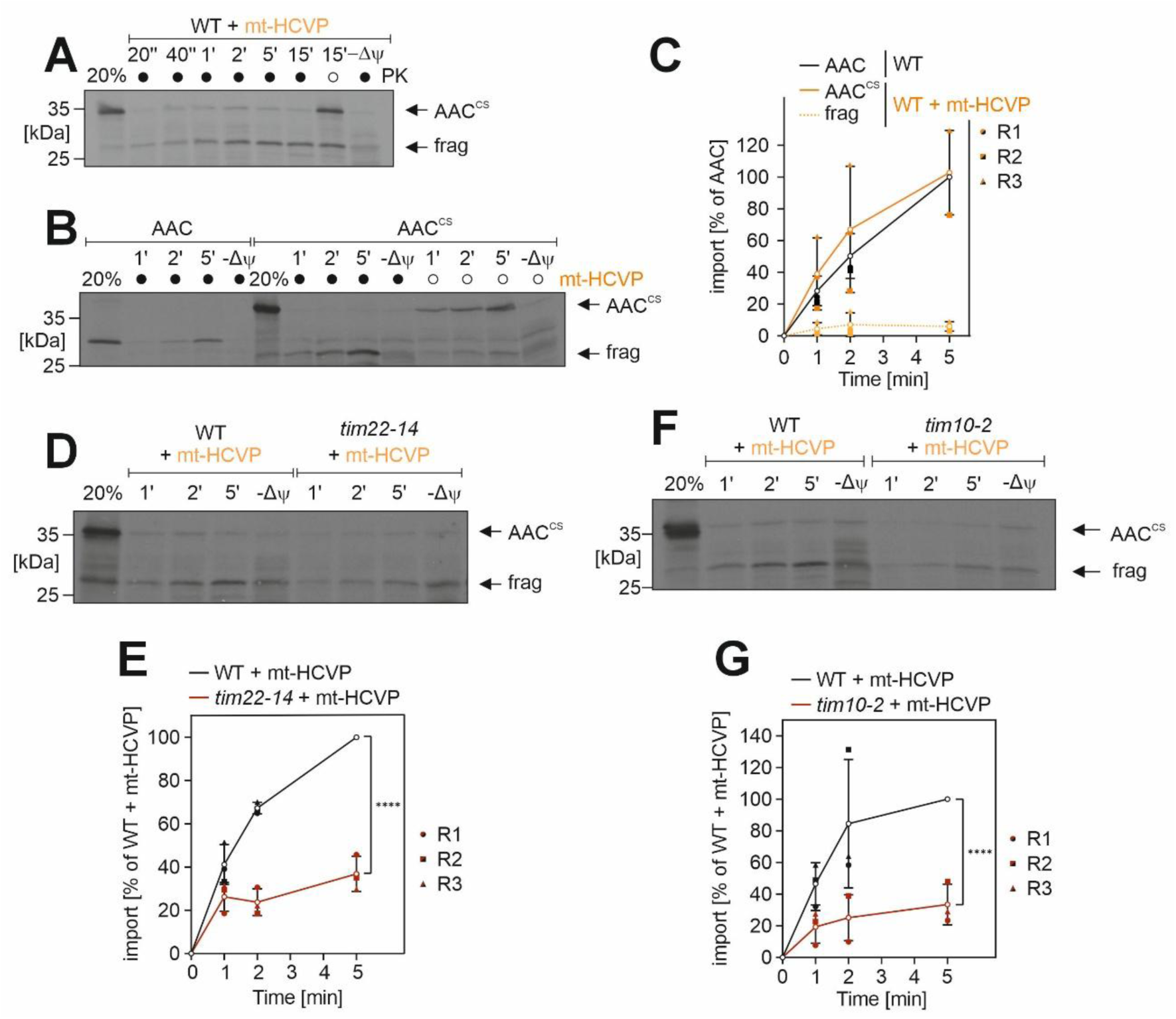
Related to Fig. 5. The insertion of carrier proteins into the inner membrane can be efficiently monitored in a protease-cleavage assay with a matrix-targeted viral protease. (A and. **B)** Radiolabeled ADP/ATP carrier AAC with or without a cleavage site (CS) was incubated at 30°C for the indicated time points with isolated mitochondria expressing matrix-targeted protease (mt-HCVP) or control mitochondria. Non-imported protein was removed by treatment with Proteinase K (PK). Some samples were treated with a mix of valinomycin, oligomycin, and antimycin A to dissipate the membrane potential (-Δψ). The 20% sample shows one-fifth of the radiolabeled protein used per time point of the import reaction. **(C)** Quantification of the in vitro cleavage assay shown in B. Shown are mean values and standard deviations of three biological (independent) replicates. frag = AAC fragment after cleavage by mt-HCVP. **(D – G)** Radiolabeled AAC with a cleavage site (CS) was incubated with isolated mitochondria from wild type, *tim22-14* or *tim10-2* cells expressing matrix-targeted hepatitis C virus protease (mt-HCVP). Samples were incubated at 35°C or 33°C for the indicated time points. Non-imported protein was removed by treatment with Proteinase K (PK). A control sample was treated with a mix of valinomycin, oligomycin, and antimycin A to dissipate the membrane potential (-Δψ). The 20% sample shows one-fifth of the radiolabeled protein used per time point of the import reaction. Mean values and standard deviations of the signals from biological (independent) replicates were quantified. Statistical analysis was performed using a two-way ANOVA with Sidak’s multiple comparison test. Statistical significance was assigned as follows: p-value < 0.0001 = ****.

## Supplemental Data

Table S1: Yeast strains used in this study

Table S2: Plasmids used in this study

Table S3: Mass spectrometry data of the Destni-Odc1 fusion protein. Related to Fig. 1F.

Table S4: GO term enrichment analysis of the Destni-Odc1 fusion protein. Related to Fig. S1J.

Table S5: Mass spectrometry data of the Aim11-HA pull-down samples. Related to Fig. 2A.

Table S6: Mass spectrometry data of the *Δaim11* and Phb1-depletion strains. Related to Fig. 3F-H.

Table S7: Heatmap data of the *Δaim11* and Phb1-depletion strains. Related to Fig. 3G.

Table S8: GO term enrichment analysis of the *Δaim11* and Phb1-depletion strains. Related to Fig. 3H.

Table S9: Complexome profiling dataset. Related to Figs. 2C and S4. Protein groups table with color-coded normalized intensity-based absolute quantification (iBAQ) values.

## Notes

### Competing Interest Statement

The authors have declared no competing interest.

